# White matter microstructure shows sex differences in late childhood: Evidence from 6,797 children

**DOI:** 10.1101/2021.08.19.456728

**Authors:** Katherine E. Lawrence, Zvart Abaryan, Emily Laltoo, Leanna M. Hernandez, Michael Gandal, James T. McCracken, Paul M. Thompson

## Abstract

Sex differences in white matter microstructure have been robustly demonstrated in the adult brain using both conventional and advanced diffusion-weighted magnetic resonance imaging (dMRI) approaches. However, sex differences in white matter microstructure prior to adulthood remain poorly understood; previous developmental work focused on conventional microstructure metrics and yielded mixed results. Here we rigorously characterized sex differences in white matter microstructure among over 6,000 children from the Adolescent Brain Cognitive Development (ABCD) Study who were between 9 and 10 years old. Microstructure was quantified using both the conventional model - diffusion tensor imaging (DTI) - and an advanced model, restriction spectrum imaging (RSI). DTI metrics included fractional anisotropy (FA) and mean, axial, and radial diffusivity (MD, AD, RD). RSI metrics included normalized isotropic, directional, and total intracellular diffusion (N0, ND, NT). We found significant and replicable sex differences in DTI or RSI microstructure metrics in every white matter region examined across the brain. Sex differences in FA were regionally specific. Across white matter regions, boys exhibited greater MD, AD, and RD than girls, on average. Girls displayed increased N0, ND, and NT compared to boys, on average, suggesting greater cell and neurite density in girls. Together, these robust and replicable findings provide an important foundation for understanding sex differences in health and disease.

## Introduction

Sex differences in white matter microstructure are consistently shown in the adult human brain (Jahanshad & Thompson, 2017; Salminen et al., 2022). Such *in vivo* neuroimaging studies have used both conventional and advanced diffusion-weighted magnetic resonance imaging (dMRI) models to extensively characterize sex differences in large-scale adult samples (Cox et al., 2016; Lawrence et al., 2021; Ritchie et al., 2018). These analyses firmly established white matter sex differences across the human brain, with some regional variability in the magnitude of such differences and the reported effect sizes indicating mean sex differences coupled with overlapping distributions in men and women (Cox et al., 2016; Lawrence et al., 2021; Ritchie et al., 2018). White matter microstructure in adulthood has also been associated with cognitive and behavioral variability, as well as a range of brain-based disorders that exhibit sex differences in their prevalence or presentation (Favre et al., 2019; Kelly et al., 2018; Meyer & Lee, 2019; Paus, Keshavan, & Giedd, 2008; Roberts, Anderson, & Husain, 2013; Salminen et al., 2022; van Velzen et al., 2020). However, we lack a complete understanding of the developmental origins of the white matter sex differences observed in adults, hindering our understanding of sex differences in health and disease.

Previous developmental work examining the impact of participant sex on white matter microstructure has used the conventional dMRI model, diffusion tensor imaging (DTI; Basser, Mattiello, & LeBihan, 1994a, 1994b), and yielded mixed findings ranging from widespread sex differences across the brain, to regionally-specific sex differences, to no significant sex differences (see Kaczkurkin, Raznahan, & Satterthwaite, 2019; Tamnes, Roalf, Goddings, & Lebel, 2018 for review). Most such studies included samples of around 100 youth or less from a range of developmental stages, such as from late childhood to late adolescence (Bava et al., 2011; Clayden et al., 2012; Herting et al., 2017; Herting, Maxwell, Irvine, & Nagel, 2012; Schmithorst, Holland, & Dardzinski, 2008; Seunarine et al., 2016). In one of the larger studies to date, Krogsrud et al. found no significant sex differences in fractional anisotropy (FA), mean diffusivity (MD), axial diffusivity (AD), or radial diffusivity (RD) at either timepoint of their longitudinal sample of 159 children aged 4 to 11 years old (Krogsrud et al., 2016). In a longitudinal study of 203 subjects scanned at 9 and 12 years old, 9-year old girls exhibited greater FA in white matter compared to their male counterparts; no sex differences were observed at 12 years of age (Brouwer et al., 2012). Lopez-Vicente et al. examined 3,031 youth aged 8 to 12 years and found significant sex differences in all white matter tracts investigated, with effect sizes indicating distributional overlap between boys and girls. On average, boys generally exhibited greater MD, AD, and RD than girls, whereas sex differences in FA were more regionally specific (Lopez-Vicente et al., 2021). As a whole, this prior work suggests sex differences in white matter microstructure during development, but the exact nature of these differences remains poorly understood.

Previous developmental studies of sex differences in white matter microstructure have used the conventional dMRI model, diffusion tensor imaging (DTI), and the resulting microstructure metrics FA, MD, AD, and RD (Kaczkurkin et al., 2019; Tamnes et al., 2018). However, the DTI model is unable to capture complex white matter architecture in the brain, such as crossing or dispersing fibers (Alexander, Lee, Lazar, & Field, 2007; Basser et al., 1994a, 1994b; Jones, 2008). The DTI metrics FA, MD, AD, and RD are also further limited by their lack of specificity in characterizing the microstructural environment. These well-established limitations of DTI are addressed by more advanced dMRI models, including the advanced model, restriction spectrum imaging (RSI). RSI models multiple underlying fiber populations per voxel, thereby allowing us to resolve complex white matter configurations, including crossing fibers (White, Leergaard, D’Arceuil, Bjaalie, & Dale, 2013; White et al., 2014; White, McDonald, et al., 2013). RSI furthermore provides microstructure measures with exhibit greater specificity than DTI. RSI is conceptually similar to the biophysical dMRI model, neurite orientation dispersion and density imaging (NODDI) (Zhang, Schneider, Wheeler-Kingshott, & Alexander, 2012). However, RSI has several strengths compared to NODDI and to signal-based diffusion models, such as mean apparent propagator MRI (MAPMRI) (Fick, Wassermann, Caruyer, & Deriche, 2016; Ozarslan et al., 2013). NODDI provides an incomplete description of intracellular diffusion, as it only models the total intracellular volume fraction. In contrast, RSI allows for greater microstructural specificity by separately reflecting isotropic and anisotropic contributions to total intracellular diffusion. RSI also allows for a more accurate estimation of fiber anisotropy when crossing fibers are perpendicular to one another, compared with the orientation dispersion measure provided by NODDI. RSI is also more biologically interpretable than signal-based models, allowing underlying neural mechanisms to be more directly addressed.

Here we thoroughly characterize sex differences in white matter microstructure in over 6,000 children from the Adolescent Brain Cognitive Development (ABCD) Study using both the conventional dMRI model, DTI, and the advanced model, RSI (Barch et al., 2018; Casey et al., 2018; Garavan et al., 2018; Hagler et al., 2019; Volkow et al., 2018). We hypothesized that the effect of sex on FA would be regionally dependent and boys would exhibit greater MD, AD, and RD than girls (Lopez-Vicente et al., 2021); the association between RSI metrics and sex was an open question due to the lack of comparable previous work. Sex differences in white matter microstructure may evolve during development. (Clayden et al., 2012; Herting et al., 2017; Schmithorst et al., 2008; Seunarine et al., 2016; Simmonds, Hallquist, Asato, & Luna, 2014 but see Krogsrud et al., 2016; Lebel & Beaulieu, 2011; Palmer, Pecheva, et al., 2021). Our analyses specifically focused on a narrow age range in late childhood, such that all participants were between 9 and 10 years old. We directly confirmed the reproducibility and robustness of all findings by using split half replication and considering a wide range of potential confounds in our analyses. To the best of our knowledge, this comprehensive study is the largest investigation to date of white matter microstructure sex differences in a developmental sample.

## Material and Methods

### Participants

The ABCD Study is a large-scale, on-going longitudinal investigation of brain development with data collected across 21 data acquisition sites (Barch et al., 2018; Casey et al., 2018; Garavan et al., 2018; Volkow et al., 2018). Participating children were recruited from schools within geographical areas that approximate the demographic diversity of the United States. Subjects completed the baseline timepoint between the ages of 9 to 10 years old. Full sampling and exclusion procedures for this epidemiologically-informed sample are described in detail elsewhere (Garavan et al., 2018). The ABCD Study protocol was approved by the centralized Institutional Review Board at the University of California, San Diego, and informed assent and consent were obtained from each participant and their legal guardian.

The current work was conducted using de-identified tabulated neuroimaging, demographic, and behavioral data from the baseline timepoint of the ABCD NIMH Data Archive (NDA) release 3.0 (ABCD DOI: 10.15154/1519007; study-specific DOI: 10.15154/1524452). Our analyses focused specifically on the baseline, late childhood timepoint of ABCD because studying sex differences prior to adolescence provides unique information on the origin of adult sex differences, compared with broader sample age ranges that include adolescents. Characterizing neural sex differences in late childhood also provides an important foundation for specifically understanding the emergence of a range of adolescent-onset neuropsychiatric disorders that exhibit clinical sex differences and display white matter alterations (Favre et al., 2019; Kelly et al., 2018; Meyer & Lee, 2019; Paus et al., 2008; Salminen et al., 2022; van Velzen et al., 2020). A complete participant flowchart for the current study is shown in **Figure S1**. Briefly, subjects were only included if they had complete nuisance covariate data (see *Statistical Analyses*) and complete dMRI data that also passed quality control (see *MRI Acquisition and Processing*) (Hagler et al., 2019). Intersex participants, or participants whose biological sex assigned at birth did not align with their parent-report gender identity, were excluded from analyses (Mueller et al., 2021). Children were also excluded if they were the sibling of another subject in the study, as statistical models run in all participants using relatedness as a random effect failed to converge; the sibling retained for analyses was selected randomly. Our final sample consisted of 6,797 children between the ages of 9 to 10 years old (48.1% female). To demonstrate the reproducibility of all findings, this final sample was split into a discovery cohort and a replication cohort. The discovery cohort included 3,399 children (47.4% female), and the replication cohort included 3,398 children (48.8% female). Descriptive statistics are presented for the discovery and replication cohorts in **Table 1**, with the reported statistical comparisons completed in R 3.6.1 using t-tests or chi-squared tests, as appropriate (R Core Team, 2016).

**Table 1.** Sample Characteristics

|  | Discovery Cohort |  | Replication Cohort |  | Discovery | Replication |
| --- | --- | --- | --- | --- | --- | --- |
|  | Male | Female | Male | Female | M vs. F | M vs. F |
| Sample Size | 1788 | 1611 | 1741 | 1657 | - | - |
| Age (years) | 9.92 ± 0.62 | 9.89 ± 0.62 | 9.93 ± 0.62 | 9.91 ± 0.61 | 0.15 | 0.46 |
| PDS Category |  |  |  |  | **< 0.001** | **< 0.001** |
| Prepubertal | 1229 | 473 | 1172 | 482 |  |  |
| Early Pubertal | 415 | 374 | 416 | 374 |  |  |
| Mid-Pubertal to Postpubertal | 88 | 711 | 99 | 742 |  |  |
| CBCL Internalizing Problems Raw | 5.06 ± 5.45 | 5.13 ± 5.61 | 5.34 ± 5.77 | 5.22 ± 5.46 | 0.74 | 0.54 |
| CBCL Externalizing Problems Raw | 4.99 ± 6.18 | 3.87 ± 5.02 | 5.17 ± 6.66 | 3.66 ± 4.92 | **< 0.001** | **< 0.001** |
| CBCL Internalizing Problems T-Score | 49.35 ± 10.74 | 47.73 ± 10.46 | 49.89 ± 10.69 | 47.98 ± 10.41 | **< 0.001** | **< 0.001** |
| CBCL Externalizing Problems T-Score | 46.39 ± 10.53 | 45.36 ± 9.65 | 46.63 ± 10.70 | 44.84 ± 9.64 | **0.003** | **< 0.001** |
| Mean Head Motion (mm) | 1.33 ± 0.49 | 1.33 ± 0.53 | 1.34 ± 0.49 | 1.33 ± 0.51 | 0.75 | 0.68 |
| Intracranial Volume (mm<sup>3</sup>) | 1.59E+06 ± 1.43E+05 | 1.45E+06 ± 1.28E+05 | 1.58E+06 ± 1.39E+05 | 1.46E+06 ± 1.28E+05 | **< 0.001** | **< 0.001** |
| Race |  |  |  |  | 0.49 | 0.50 |
| Asian | 38 | 39 | 38 | 40 |  |  |
| Black | 253 | 243 | 253 | 260 |  |  |
| White | 1214 | 1054 | 1155 | 1058 |  |  |
| Other/Mixed | 283 | 275 | 295 | 299 |  |  |
| Ethnicity |  |  |  |  | 0.32 | 0.84 |
| Hispanic or Latino/Latina | 342 | 330 | 361 | 339 |  |  |
| Not Hispanic or Latino/Latina | 1446 | 1281 | 1380 | 1318 |  |  |
| Household Income |  |  |  |  | 0.77 | 0.50 |
| Less than \$50k | 512 | 477 | 518 | 503 | | |
| Between \$50k - \$100k | 508 | 443 | 497 | 495 | | |
| More than \$100k | 768 | 691 | 726 | 659 | | |
| Parental Education |  |  |  |  | 0.36 | 0.74 |
| Less than HS Diploma | 57 | 71 | 55 | 60 |  |  |
| HS Diploma / GED | 140 | 131 | 160 | 133 |  |  |
| Some College | 489 | 414 | 454 | 430 |  |  |
| Bachelor Degree | 467 | 420 | 463 | 443 |  |  |
| Post-Graduate Degree | 635 | 575 | 609 | 591 |  |  |

Descriptive statistics are presented as mean ± standard deviation for continuous variables and as subject counts for discrete variables. PDS = Pubertal Development Scale. CBCL = Child Behavior Checklist. HS = High School. GED = General Education Development test. PDS data was missing for 109 subjects in the discovery cohort and 113 subjects in the replication cohort. CBCL data was missing for 1 subject in the replication cohort

### MRI Acquisition and Processing

T1-weighted structural MRI and dMRI scans were acquired as described previously (Casey et al., 2018). Briefly, scans were acquired using a harmonized acquisition protocol across 21 ABCD sites on Siemens Prisma, Philips, or GE 750 3-tesla scanners. The 3D T1-weighted magnetization-prepared rapid acquisition gradient echo (MPRAGE) structural MRI scan had isotropic voxel dimensions of 1 mm and was collected without multi-band acceleration. Diffusion-weighted MRI scans consisted of 96 diffusion encoding directions across four diffusion-weighted shells: 6 directions at *b* = 500 s/mm^2^, 15 directions at *b* = 1000 s/mm^2^, 15 directions at *b* = 2000 s/mm^2^, and 60 directions at *b* = 3000 s/mm^2^. Voxel dimensions were 1.7 mm isotropic and a multi-band acceleration factor of 3 was used.

Processing steps for the T1-weighted structural MRI and dMRI scans are detailed extensively elsewhere (Hagler et al., 2019). In brief, processing was completed using the Multi-Modal Processing Stream (MMPS; https://www.nitrc.org/projects/abcd_study), a software package that calls on in-house ABCD scripts, as well as the publicly available neuroimaging tools FreeSurfer (Fischl et al., 2002), Analysis of Functional NeuroImages (AFNI; Cox, 1996), and FMRIB Software Library (FSL; Jenkinson, Beckmann, Behrens, Woolrich, & Smith, 2012; Smith et al., 2004). T1-weighted images underwent correction for gradient non-linearity distortions using scanner-specific, nonlinear transformations provided by the MRI scanner manufacturers (Jovicich et al., 2006; Wald, Schmitt, & Dale, 2001) and were subsequently registered to standard space (Friston et al., 1995). Preprocessing for the diffusion-weighted MRI scans included eddy current correction, gradient distortion correction using pairs of reverse-phase encoded *b* = 0 s/mm^2^ dMRI images, and registration to the T1-weighted scan. To reduce the potential impact of head motion, motion-contaminated slices were censored prior to diffusion model fitting (Hagler et al., 2019). Processed dMRI images were also visually inspected for residual motion artifacts (Hagler et al., 2019).

Two reconstruction models were fit to the dMRI data: the conventional model, DTI, and the advanced model, RSI (Basser et al., 1994a, 1994b; Hagler et al., 2019; Jones, 2008; White, Leergaard, et al., 2013; White et al., 2014; White, McDonald, et al., 2013). DTI models a single fiber orientation per voxel by fitting a single diffusion tensor, or ellipsoid, for each voxel in the brain (Basser et al., 1994a, 1994b). The DTI model was fit using diffusion-weighted shells equal to or less than 1000 s/mm^2^ (inner shell DTI) and, separately, using all collected gradient strengths (full shell DTI) (Hagler et al., 2019). To improve comparability with previous work, our analyses here focus on the inner shell DTI measures (Kaczkurkin et al., 2019; Tamnes et al., 2018). Microstructure metrics derived from DTI included FA, MD, AD, and RD. FA reflects the degree of diffusion anisotropy, and MD represents the average diffusivity in all directions. AD reflects the diffusion parallel to the primary diffusion axis, and RD captures the diffusion perpendicular to the primary diffusion axis. Conventionally, but with several caveats, higher FA is considered to reflect greater white matter integrity or a reduction in crossing fibers (Alexander et al., 2007; Thomason & Thompson, 2011). Higher MD is conventionally thought to indicate increased extracellular volumes or decreased cellular density, lower AD is conventionally believed to reflect axonal pruning or axonal damage, and increased RD is conventionally related to less myelination or the presence of myelin injury (Alexander et al., 2007; Song et al., 2003; Song et al., 2002). However, the DTI model has well-established limitations regarding its inability to resolve complex fiber configurations and the non-specificity of its microstructure metrics (Alexander et al., 2007; Basser et al., 1994a, 1994b; Jones, 2008).

RSI is an advanced diffusion model that allows for the resolution of complex white matter architecture, including crossing fibers, and provides microstructure indices with greater specificity than DTI (White, Leergaard, et al., 2013; White et al., 2014; White, McDonald, et al., 2013). RSI separately models restricted (intracellular) and hindered (extracellular) diffusion in each voxel using fourth order spherical harmonic functions, allowing for the modeling of multiple diffusion orientations within each voxel. Our analyses here focus on RSI metrics derived from intracellular diffusion, as these may provide more intuitive interpretations than measures calculated from extracellular diffusion by virtue of more directly reflecting diffusion within cell bodies and neurites (White, Leergaard, et al., 2013; White et al., 2014; White, McDonald, et al., 2013). Here we analyzed the following intracellular microstructure indices from RSI (Hagler et al., 2019): normalized isotropic (N0), normalized directional (ND) and normalized total (NT) intracellular diffusion. Each RSI metric is defined as the Euclidean norm of the corresponding model coefficients divided by the norm of all model coefficients and is a unitless measure that ranges from 0 to 1. N0 is derived from the 0^th^ order spherical harmonic coefficient and reflects isotropic intracellular diffusion. ND is calculated from the norm of the 2^nd^ and 4^th^ order spherical harmonic coefficients and captures oriented intracellular diffusion. NT is derived from the norm of the 0^th^, 2^nd^, and 4^th^ order spherical harmonic coefficients and reflects the overall intracellular diffusion. A number of biological processes may contribute to the intracellular signal fraction, including variability in myelination, astrocytes, and microglia. In addition to the potential impact of these processes, higher N0 may reflect increased cell density, higher ND may relate to increased neurite density, and greater NT values may indicate an overall larger intracellular space. These RSI indices thus provide distinct and complementary microstructural information by separately reflecting neurite density and cell density; together, these metrics allow for a more comprehensive characterization of the underlying neurobiology.

Major white matter tracts across the brain were labeled using the well-established probabilistic fiber tract atlas, AtlasTrack (Gustavson et al., 2019; Hagler et al., 2009; Hagler et al., 2019; Kaestner et al., 2020; Reyes et al., 2018; Ursache et al., 2016). More specifically, subjects’ T1-weighted MRI images were first registered nonlinearly to the atlas. Diffusion orientations were then compared between each subjects’ dMRI scan and the atlas to refine *a priori* tract locations and to personalize each white matter region of interest (ROI), including minimizing the contribution of atlas-inconsistent regions. Voxels which were identified by FreeSurfer as containing primarily gray matter or cerebrospinal fluid were also excluded.

Mean values for each DTI and RSI metric were extracted for each white matter fiber tract ROI, and the following bilateral ROIs were examined in the current study (**Figure S2, Table S1**): all white matter fibers, anterior thalamic radiation (ATR), corpus callosum (CC), cingulate cingulum (CGC), parahippocampal cingulum (CGH), corticospinal/pyramidal tract (CST), *forceps major* (Fmaj), *forceps minor* (Fmin), fornix (FX), fornix excluding the fimbria (FXcut), inferior fronto-occipital fasciculus (IFO), inferior frontal superior frontal cortex (IFSFC), inferior longitudinal fasciculus (ILF), superior corticostriate (SCS), frontal superior corticostriate (fSCS), parietal superior corticostriate (pSCS), striatal inferior frontal cortex (SIFC), superior longitudinal fasciculus (SLF), parietal superior longitudinal fasciculus (pSLF), temporal superior longitudinal fasiculus (tSLF), and uncinate fasciculus (UNC).

Processed dMRI data were visually inspected for quality control purposes, as described previously (Hagler et al., 2019). Briefly, each subject’s co-registered T1-weighted and ND images were assessed for residual distortion, registration quality, image quality (including residual motion artifacts), and automatic white matter tract segmentation. Participants’ dMRI data was only included in the current study if they passed quality control on all four dimensions assessed.

### Statistical Analyses

Linear mixed-effects models were used to examine microstructural sex differences in white matter fiber tract ROIs across the brain. These models were used as they allow for nested data structures while appropriately modeling non-independent data points, such as participants nested within MRI scanners; linear mixed-effects models have also been used extensively in previously published multi-scanner neuroimaging studies, including prior work in ABCD (e.g., Bohon & Welch, 2021; Hernandez et al., 2022; Pagliaccio, Durham, Fitzgerald, & Marsh, 2021; Palmer, Pecheva, et al., 2021; Palmer, Zhao, et al., 2021; Piccolo et al., 2016; Rapuano et al., 2020). All regressions for the current study were completed in R 3.6.1 using the *lme4* package and included the following standard nuisance covariates as fixed effects: age, household income, parental education, race, and ethnicity; MRI scanner was modeled as a random effect in all analyses to adjust for scanner-related variability by modeling the hierarchical structure of our multi-scanner dataset (Bohon & Welch, 2021; Hernandez et al., 2022; Pagliaccio et al., 2021; Palmer, Pecheva, et al., 2021; Palmer, Zhao, et al., 2021; Rapuano et al., 2020). Results were considered significant and replicable if they survived a 5% false discovery rate (FDR) applied across the number of ROIs in the discovery cohort (*q*<0.05) and demonstrated *p*<0.05 in the replication cohort (Benjamini & Hochberg, 1995; Hernandez et al., 2022). Effect sizes are reported for all analyses as the magnitude of the standardized regression coefficients (standardized betas; βs), reflecting that group comparisons were completed as regressions to allow for the inclusion of nuisance covariates; reporting effect sizes as standardized betas is consistent with prior work and also allows for greater comparability with previously published ABCD studies (e.g., Cai, Elsayed, & Barch, 2021; Cox et al., 2016; Hernandez et al., 2022; Lawrence et al., 2020; Lopez-Vicente et al., 2021; Pagliaccio et al., 2021; Palmer, Zhao, et al., 2021).

Primary analyses characterized the effect of sex on inner shell DTI metrics and intracellular RSI metrics in the bilateral fiber tract ROIs (**Table S1**); unless otherwise specified, all subsequent references to DTI correspond to inner shell DTI, and all subsequent references to RSI correspond to intracellular RSI. A number of analyses were completed to establish the robustness of our primary findings. First, we examined sex differences separately in the left and right hemispheres. Second, we included pubertal development as a covariate to statistically control for known sex differences in pubertal maturation among this age group (Herting et al., 2020). Third, we repeated primary analyses when covarying for raw dimensional externalizing or internalizing problem scores from the parent-report CBCL. Fourth, we statistically controlled for mean head motion (Yendiki, Koldewyn, Kakunoori, Kanwisher, & Fischl, 2014). Fifth, intracranial volume was included as a nuisance covariate, as some recent work has posited that sex differences in brain connectivity are driven by known differences in brain size between males and females (Eliot, Ahmed, Khan, & Patel, 2021; Salminen et al., 2022). Sixth, we statistically covaried for all supplemental regressors within a single analysis, including pubertal development, externalizing problems, internalizing problems, head motion, and intracranial volume. Lastly, we used an alternative statistical approach to control for scanner-related variability, ComBat (Fortin et al., 2017); ComBat is an advanced harmonization approach that uses an empirical Bayes framework to correct for scanner effects and has previously been used in a range of multi-site neuroimaging studies (e.g., Nir et al., 2019; Radua et al., 2020; Thomopoulos et al., 2021; Zavaliangos-Petropulu et al., 2019). For completeness, we also repeated our primary analyses when combining the discovery and replication cohorts into a single cohort, and we investigated sex differences in full shell DTI and extracellular RSI indices.

Follow-up analyses expanded on our primary DTI and RSI findings by directly assessing the relative sensitivity of these two models to sex differences. Specifically, we statistically compared the effect size of the most sensitive DTI metric with that of the most sensitive RSI metric for each fiber tract ROI. In line with all other analyses, results were only deemed significant and replicable if they survived FDR correction in the discovery cohort (*q*<0.05) and demonstrated *p*<0.05 in the replication cohort (Hernandez et al., 2022).

## Results

Participant demographic information is presented for the discovery and replication cohorts in **Table 1**. Boys and girls were matched on all core demographic variables, including age, household income, parental education, race, and ethnicity (**Table 1**). As expected, significant sex differences were observed in pubertal development, externalizing behaviors, and intracranial volume (**Table 1**). Boys were less pubertally advanced than girls, as assessed by the parent-report Pubertal Development Scale (PDS; Petersen, Crockett, Richards, & Boxer, 1988). Boys also exhibited higher measures of externalizing behaviors than girls, as quantified by raw externalizing problem scores from the parent-report Child Behavior Checklist (CBCL; Achenbach, 2009). Intracranial volume was significantly larger among boys than girls. No significant differences were observed between boys and girls in raw internalizing problem scores from the parent-report CBCL or in mean head motion (**Table 1**).

White matter sex differences were observed across the brain, with significant and replicable sex differences found in DTI or RSI microstructure metrics for every bilateral white matter tract ROI examined (**Figure 1A, Table S2-S8**). The directionality of sex differences in FA varied across white matter ROIs. FA was higher in boys than girls in limbic white matter tract ROIs (CGC, CGH, FX, FXcut) and thalamic projections (ATR) (discovery βs=0.08-0.21; replication βs=0.09-0.20), on average. In contrast, greater FA was observed among girls in association tract ROIs (IFO, ILF, tSLF, IFSFC, UNC), corticostriatal projections (SCS, fSCS, pSCS, SIFC), and across all white matter fibers on average, compared to boys (discovery βs=0.08-0.25; replication βs=0.07-0.27). Sex differences were highly consistent across ROIs for all other examined microstructure indices: MD, AD, RD, N0, ND, and NT. Boys typically exhibited greater MD, AD, and RD than girls, on average, indicating higher diffusivity in boys (discovery βs=0.12-0.49; replication βs=0.07-0.43). Girls generally displayed higher N0, ND, and NT relative to boys, suggesting greater cell and neurite density in girls (discovery βs=0.08-0.45; replication βs=0.07-0.42). Across microstructure measures, the most substantial sex differences were observed in association tract ROIs (IFO, ILF, tSLF, IFSFC) and superior corticostriatal projections (SCS, fSCS, pSCS). Sex differences were the least pronounced, albeit still significant and replicable, in limbic white matter tract ROIs (FX, FXcut), commissural tract ROIs (Fmin, Fmaj), and sensorimotor and thalamic projections (CST, ATR).

**Figure 1.**
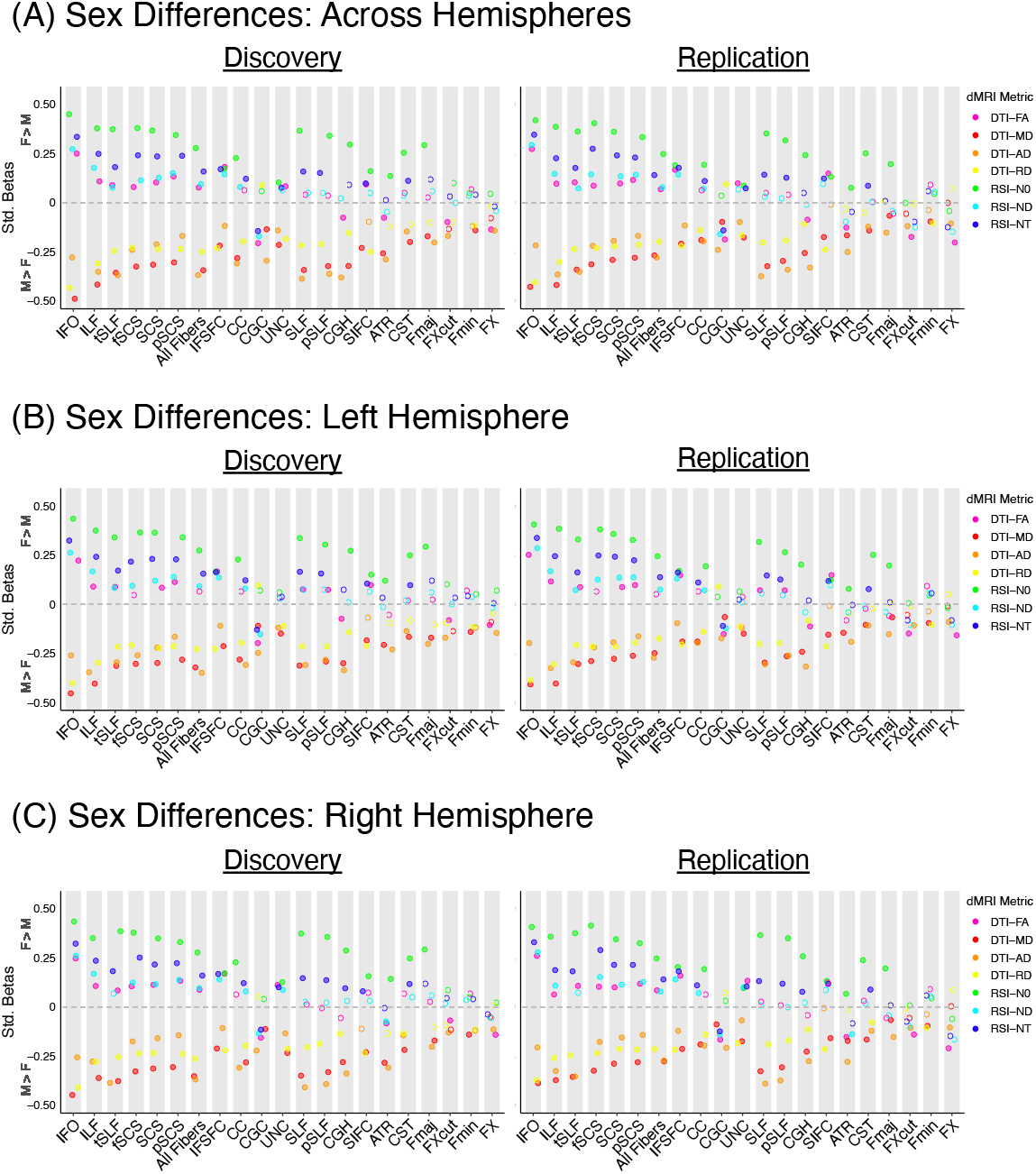
Sex differences in white matter microstructure. The effect size, significance, and replicability of sex differences in DTI and RSI metrics are depicted separately for the discovery cohort (left) and replication cohort (right) for (A) bilateral white matter tract ROIs, (B) left hemisphere white matter tract ROIs, and (C) right hemisphere white matter tract ROIs; commissural tracts are included in the left and right hemisphere graphs for completeness only. Positive standardized betas indicate Female > Male, and negative standardized betas indicate Male > Female. DTI metrics are depicted in warm colors, and RSI metrics in cool colors. Filled circles indicate the association was both significant in the discovery cohort after FDR correction across the number of ROIs (*q*<0.05) and demonstrated *p*<0.05 in the replication cohort. *Diffusion-weighted MRI abbreviations*: ROI = region of interest, dMRI = diffusion-weighted MRI, DTI = diffusion tensor imaging, FA = fractional anisotropy, MD = mean diffusivity, AD = axial diffusivity, RD = radial diffusivity, RSI = restriction spectrum imaging, N0 = normalized isotropic, ND = normalized directional, NT = normalized total. *White matter tract ROI abbreviations*: ATR = anterior thalamic radiation, CC = corpus callosum, CGC = cingulum (cingulate), CGH = cingulum (parahippocampal), CST = corticospinal/pyramidal tract, Fmaj = *forceps major*, Fmin = *forceps minor*, fSCS = superior corticostriate (frontal cortex), FX = fornix, FXcut = fornix (excluding fimbria), IFO = inferior fronto-occipital fasciculus, IFSFC = inferior frontal superior frontal cortex, ILF = inferior longitudinal fasciculus, pSCS = superior corticostriate (parietal cortex), pSLF = superior longitudinal fasciculus (parietal), SCS = superior corticostriate, SIFC = striatal inferior frontal cortex, SLF = superior longitudinal fasciculus, tSLF = superior longitudinal fasciculus (temporal), UNC = uncinate fasciculus.

Follow-up analyses considering sex differences separately in the left and right hemisphere yielded highly similar results to our primary analyses examining bilateral white matter tract ROIs (**Figure 1B-C**, discovery βs=0.05-0.45; replication βs=0.05-0.41). Results were also largely consistent when controlling for pubertal development (**Figure 2**, discovery βs=0.08-0.50; replication βs=0.06-0.40); a small number of ROI and dMRI metric combinations no longer exhibited significant and replicable sex effects, and sex differences were now also observed for FA in commissural tract ROIs (CC, Fmin). Follow-up analyses including dimensional externalizing or internalizing problems as a nuisance covariate provided very similar results to our primary analyses (**Figure 3A-B**, discovery βs=0.05-0.49; replication βs = 0.06-0.43). In sum, our sex difference findings were highly consistent across hemispheres, as well as when statistically controlling for pubertal development, externalizing problems, and internalizing problems.

**Figure 2.**
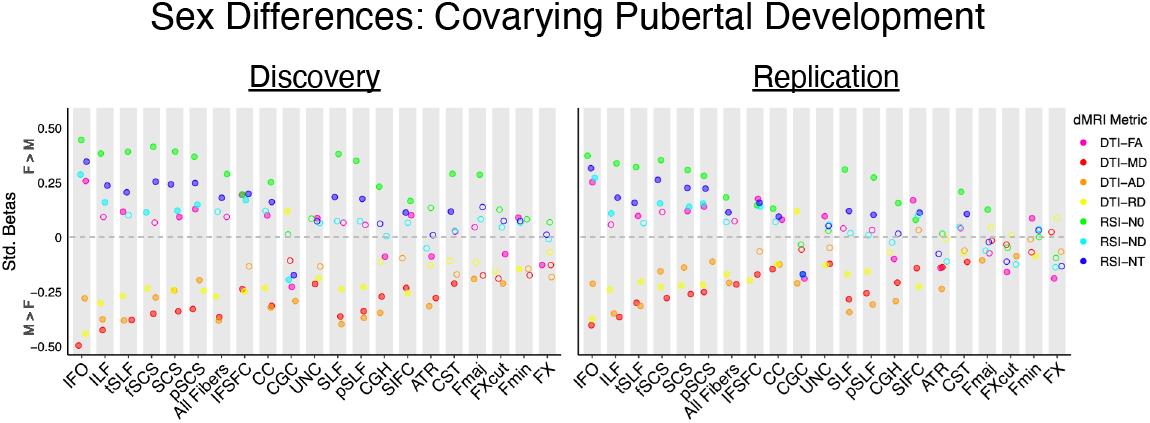
Sex differences in white matter microstructure, controlling for pubertal development. The effect size, significance, and replicability of sex differences in DTI and RSI metrics are depicted separately for the discovery cohort (left) and replication cohort (right) for bilateral white matter tract ROIs. Positive standardized betas indicate Female > Male, and negative standardized betas indicate Male > Female. DTI metrics are depicted in warm colors, and RSI metrics in cool colors. Filled circles indicate the association was both significant in the discovery cohort after FDR correction across the number of ROIs (*q*<0.05) and demonstrated *p*<0.05 in the replication cohort. *Diffusion-weighted MRI abbreviations*: ROI = region of interest, dMRI = diffusion-weighted MRI, DTI = diffusion tensor imaging, FA = fractional anisotropy, MD = mean diffusivity, AD = axial diffusivity, RD = radial diffusivity, RSI = restriction spectrum imaging, N0 = normalized isotropic, ND = normalized directional, NT = normalized total. *White matter tract ROI abbreviations*: ATR = anterior thalamic radiation, CC = corpus callosum, CGC = cingulum (cingulate), CGH = cingulum (parahippocampal), CST = corticospinal/pyramidal tract, Fmaj = *forceps major*, Fmin = *forceps minor*, fSCS = superior corticostriate (frontal cortex), FX = fornix, FXcut = fornix (excluding fimbria), IFO = inferior fronto-occipital fasciculus, IFSFC = inferior frontal superior frontal cortex, ILF = inferior longitudinal fasciculus, pSCS = superior corticostriate (parietal cortex), pSLF = superior longitudinal fasciculus (parietal), SCS = superior corticostriate, SIFC = striatal inferior frontal cortex, SLF = superior longitudinal fasciculus, tSLF = superior longitudinal fasciculus (temporal), UNC = uncinate fasciculus.

**Figure 3.**
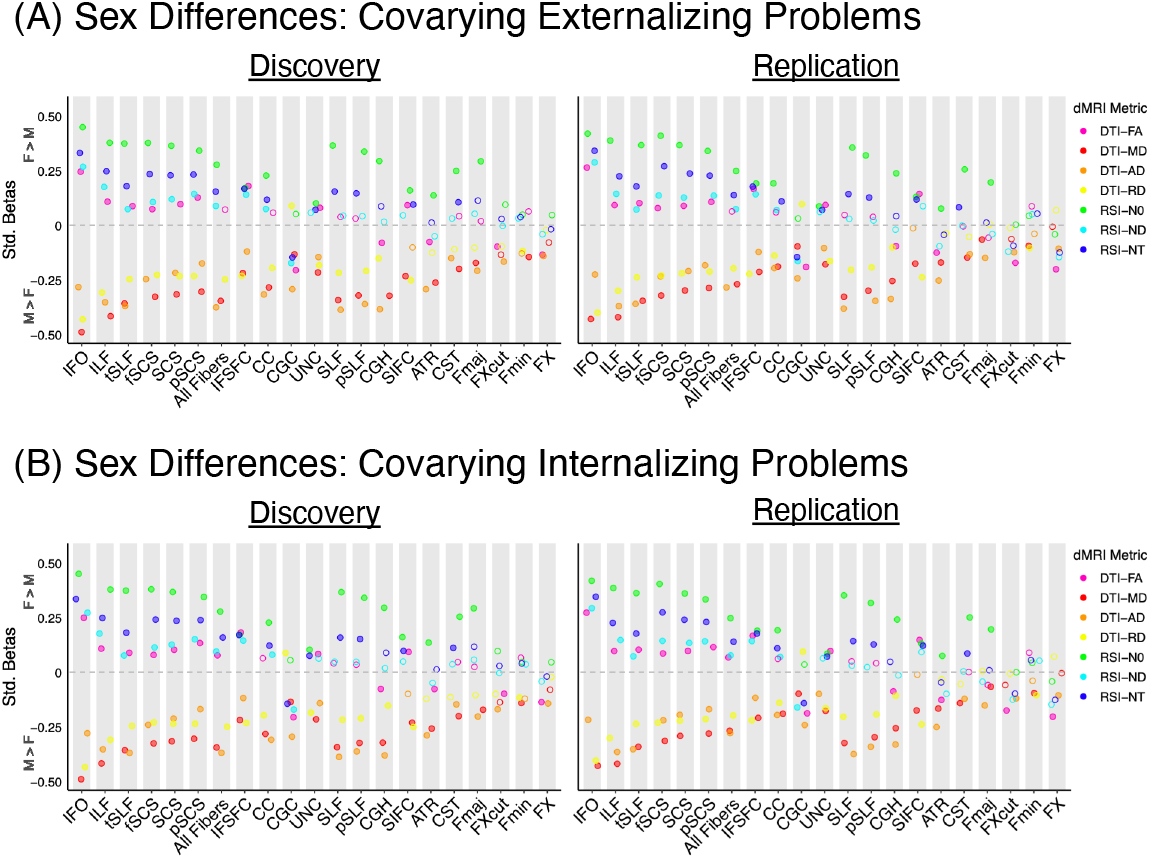
Sex differences in white matter microstructure, controlling for dimensional externalizing and internalizing problems. The effect size, significance, and replicability of sex differences in DTI and RSI metrics are depicted separately for the discovery cohort (left) and replication cohort (right) for bilateral white matter tract ROIs when controlling for (A) dimensional externalizing problems and (B) dimensional internalizing problems. Positive standardized betas indicate Female > Male, and negative standardized betas indicate Male > Female. DTI metrics are depicted in warm colors, and RSI metrics in cool colors. Filled circles indicate the association was both significant in the discovery cohort after FDR correction across the number of ROIs (*q*<0.05) and demonstrated *p*<0.05 in the replication cohort. *Diffusion-weighted MRI abbreviations*: ROI = region of interest, dMRI = diffusion-weighted MRI, DTI = diffusion tensor imaging, FA = fractional anisotropy, MD = mean diffusivity, AD = axial diffusivity, RD = radial diffusivity, RSI = restriction spectrum imaging, N0 = normalized isotropic, ND = normalized directional, NT = normalized total. *White matter tract ROI abbreviations*: ATR = anterior thalamic radiation, CC = corpus callosum, CGC = cingulum (cingulate), CGH = cingulum (parahippocampal), CST = corticospinal/pyramidal tract, Fmaj = *forceps major*, Fmin = *forceps minor*, fSCS = superior corticostriate (frontal cortex), FX = fornix, FXcut = fornix (excluding fimbria), IFO = inferior fronto-occipital fasciculus, IFSFC = inferior frontal superior frontal cortex, ILF = inferior longitudinal fasciculus, pSCS = superior corticostriate (parietal cortex), pSLF = superior longitudinal fasciculus (parietal), SCS = superior corticostriate, SIFC = striatal inferior frontal cortex, SLF = superior longitudinal fasciculus, tSLF = superior longitudinal fasciculus (temporal), UNC = uncinate fasciculus.

To further assess the robustness of our sex difference results, we repeated analyses when controlling for head motion, intracranial volume, and all supplemental nuisance covariates at once; we also repeated analyses when using an alternate approach to control for scanner-related variability, ComBat. Results were highly similar when covarying for head motion (**Figure S3**, discovery βs=0.07-0.49; replication βs=0.06-0.43). Patterns of sex differences were also largely comparable when controlling for intracranial volume (**Figure S4**, discovery βs=0.07-0.42; replication βs=0.05-0.43). Some ROI and dMRI metric combinations no longer exhibited significant and replicable sex effects, or vice versa; a small number of ROI and DTI metrics, but not RSI metrics, exhibited reversed directionality of their findings (**Figure S4**). When statistically controlling for all supplemental covariates within a single analyses – including pubertal development, externalizing problems, internalizing problems, head motion, and intracranial volume – the effect of participant sex was highly similar to our results when covarying for intracranial volume alone (**Figure S5**, discovery βs=0.12-0.38; replication βs=0.08-0.38); significant and replicable sex differences were no longer observed for some ROI and dMRI measure combinations. ComBat-based scanner harmonization yielded highly similar results to those in our primary analyses, which included MRI scanner as a random effect (**Figure S6**, discovery βs=0.07-0.50; replication βs=0.07-0.45). As a whole, our patterns of observed sex differences were largely consistent when controlling for head motion, intracranial volume, and all supplemental covariates at once, as well as when using ComBat to harmonize scanner.

For completeness, primary analyses were repeated when combining the discovery and replication cohorts into a single cohort, and when examining full shell DTI and extracellular RSI metrics. Patterns of sex differences remained consistent in the full sample, with additional significant differences observed in DTI and RSI indices for some ROIs (**Figure S7**, βs=0.04-0.46). Similar to our primary analyses characterizing inner shell DTI and intracellular RSI measures, full shell DTI and extracellular RSI indices indicated greater white matter diffusivity in boys than girls, as well as increased cell and neurite density in girls (**Figure S8**, discovery βs=0.05-0.37; replication βs=0.06-0.38). Together, these results further support widespread, significant sex differences in DTI and RSI measures of white matter microstructure.

Lastly, we expanded on our primary analyses by considering the relative sensitivity of the analyzed DTI and RSI metrics to sex differences. Among DTI metrics, MD or AD generally detected sex differences the most sensitively. N0 typically exhibited the greatest sensitivity among the RSI metrics. When directly contrasting the relative sensitivity of DTI and RSI measures to sex effects, minimal differences were found. The only significant and replicable difference was observed in the ATR, such that the DTI metric AD was more sensitive to sex effects than the RSI metric N0 (discovery FDR=0.015; replication *p*=8.5E-05).

## Discussion

Here we rigorously characterized sex differences in white matter microstructure in the largest developmental sample to date using both the conventional dMRI model, DTI, and the advanced model, RSI. Previous work in adults has robustly established white matter sex differences in both conventional and advanced microstructure measures, with some regional variability in the magnitude of such differences (Jahanshad & Thompson, 2017; Salminen et al., 2022). However, potential sex differences in white matter microstructure prior to adulthood have remained poorly understood. Prior developmental work only included conventional microstructure metrics and yielded mixed results, which varied from widespread sex differences to no significant sex differences (Kaczkurkin et al., 2019; Tamnes et al., 2018).

In our sample of over 6,000 children between the ages of 9 to 10 years old, we found significant and replicable sex differences in DTI and RSI microstructure measures across the brain. Sex differences in FA were regionally specific, whereas sex differences in MD, AD, and RD were generally consistent across white matter regions, such that boys exhibited greater diffusivity than girls, on average. Girls typically displayed increased N0, ND, and NT compared to boys, on average, suggesting increased cell and neurite density in girls. Given that our results remained consistent when statistically controlling for puberty, these observed microstructure sex differences in late childhood may be due to the impact of X chromosome genes and/or the organizational effects of intrauterine testosterone (Lentini, Kasahara, Arver, & Savic, 2013; Mallard et al., 2021; Salminen et al., 2022).

With regard to the regional distribution of sex differences, differences were most pronounced in association and superior corticostriatal white matter tract ROIs. Association fibers connect distributed cortical regions within the same hemisphere, and corticostriatal tracts connect cortical areas across the brain with the striatum (Buyanova & Arsalidou, 2021; Hagler et al., 2009). The impact of participant sex was the least notable in commissural and limbic (fornix) white matter tract ROIs, as well as sensorimotor (corticospinal/pyramidal tract) and thalamic (anterior thalamic radiation) projection tract ROIs. Commissural tracts connect the left and right hemispheres, and the fornix connects areas involved in memory (Buyanova & Arsalidou, 2021; Catani & Thiebaut de Schotten, 2008; Hagler et al., 2009). The corticospinal/pyramidal tract connects the motor cortex to the spinal cord, and the anterior thalamic radiation connects the thalamus with the frontal cortex (Buyanova & Arsalidou, 2021; Catani & Thiebaut de Schotten, 2008; Hagler et al., 2009). As association tracts generally exhibit more prolonged development than commissural and projection tracts, our results suggest that sex differences in late childhood may be most substantial in fibers with protracted developmental trajectories (Tamnes et al., 2018).

Sex differences in white matter were assessed in our sample by applying both the conventional microstructural model, DTI, and the advanced microstructural model, RSI. When directly contrasting the sensitivity of these two models to sex differences, the two approaches were largely comparable. However, the DTI model has well-established limitations regarding its inability to resolve complex white matter architecture and the non-specificity of its microstructure metrics (Alexander et al., 2007; Basser et al., 1994a, 1994b; Jones, 2008). RSI, in comparison, can resolve complex fiber configurations and provides more refined microstructure measures, offering greater insight into the underlying neurobiology than DTI (White, Leergaard, et al., 2013; White et al., 2014; White, McDonald, et al., 2013). Prior work has together indicated that the relative sensitivity of conventional and advanced dMRI metrics may also depend on the specific neurobiology underlying the scientific question of interest, allowing for the possibility that DTI and RSI may exhibit significantly different sensitivity to effects beyond participant sex (Lawrence et al., 2021; Nir et al., 2019; Pines et al., 2020).

As a whole, our findings expand on prior developmental studies that reported conflicting results when examining sex differences in FA, MD, AD, or RD in samples of around 200 youth or less (Bava et al., 2011; Brouwer et al., 2012; Clayden et al., 2012; Herting et al., 2017; Herting et al., 2012; Krogsrud et al., 2016; Schmithorst et al., 2008; Seunarine et al., 2016). Supporting the importance of large sample sizes in stabilizing results (Button et al., 2013; Marek et al., 2020), our results are generally consistent with the largest previously published study of developmental sex effects in FA, MD, AD, and RD, which assessed 3,031 youth aged 8 to 12 years old (Lopez-Vicente et al., 2021). Intriguingly, our sex difference findings here contrast with those of previous adult sex difference studies, which examined large-scale samples ranging from 3,513 to 15,628 adults (Cox et al., 2016; Lawrence et al., 2021; Ritchie et al., 2018). We found here that late childhood sex differences in FA were regionally specific, but sex differences in MD, AD, and RD were generally consistent across white matter regions. The opposite pattern is observed in adults: sex differences in FA are generally consistent across tracts in adulthood, but sex differences in MD, AD, and RD are regionally specific (Cox et al., 2016; Lawrence et al., 2021; Ritchie et al., 2018). Across microstructure metrics, the regional distribution of sex differences also differs between our late childhood sample and previously analyzed adult samples. Sex differences in adulthood are particularly pronounced in sensorimotor tracts (Cox et al., 2016; Ritchie et al., 2018). In contrast, our findings here indicate that late childhood sex differences are especially small in sensorimotor tracts. Together, these disparate patterns of findings between late childhood and adulthood suggest sex-specific maturational trajectories during adolescence. Some prior work including adolescent samples has likewise suggested that sex differences may evolve over development (Clayden et al., 2012; Herting et al., 2017; Schmithorst et al., 2008; Seunarine et al., 2016; Simmonds et al., 2014 but see Lebel & Beaulieu, 2011), although sex-specific trajectories may not arise until after early adolescence (Palmer, Pecheva, et al., 2021). Sex differences in maturational trajectories between childhood and adulthood may be driven by pubertal development and the associated changes in sex hormones during adolescence (Crone & Dahl, 2012). Indeed, previous work in humans has suggested an association between sex hormones, such as estrogen or testosterone, and white matter microstructure (Herting et al., 2012; Nabulsi et al., 2020; van Hemmen et al., 2017). The impact of estrogen, progesterone, and testosterone on white matter has also been confirmed in animal studies, which have demonstrated that such hormones directly impact processes such as oligodendrocyte proliferation, maturation, and cell death (Cerghet, Skoff, Swamydas, & Bessert, 2009; Schumacher et al., 2012).

The current study has a number of important strengths, including the unprecedented sample size, the use of both conventional and advanced microstructure models, the narrow age range of the sample, and the demonstrated reproducibility and robustness of our results. Our findings provide an important foundation for future work dissecting the mechanisms driving sex differences in white matter microstructure, including the potential impact of pubertal development, sex hormones, genetic factors, and environmental contributors (Herting et al., 2017; Herting et al., 2012; Mallard et al., 2021; Nabulsi et al., 2020; Salminen et al., 2022). These findings also lay the groundwork for future studies examining how white matter sex differences may relate to individual variability in brain function and behavior, including how such neural differences may relate to clinical sex differences in adolescent-onset neuropsychiatric disorders (Meyer & Lee, 2019; Paus et al., 2008; Salminen et al., 2022; van Eijk et al., 2021). Future work should address the limitations of the current study by examining white matter sex differences in younger samples and charting such differences longitudinally from childhood through adulthood, including assessing whether potential sex differences in maturational trajectories are mediated by pubertal hormones (Tamnes et al., 2018). Additionally, further animal and postmortem studies are needed to more precisely delineate the exact neurobiology underpinning white matter sex differences during development.

In conclusion, we thoroughly characterized white matter sex differences in late childhood using both conventional and advanced microstructure measures. Our results demonstrate replicable and robust sex differences in white matter microstructure across the brain in the largest developmental sample to date. These findings provide an important foundation for understanding sex differences in health and disease.

## Acknowledgments

This research was conducted using data obtained from the ABCD NIMH Data Archive (NDA) release 3.0 (DOI: 10.15154/1519007).

## Notes

**Funding Statement:** This work was supported by the National Institute of Mental Health (grant numbers R01MH116147 to P.M.T., F32MH122057 to K.E.L., and K00MH119663 to L.M.H.).

**Conflict of Interest Disclosure:** P.M.T. reports a research grant from Biogen, Inc. for research unrelated to this manuscript. J.T.M. reports consultant income from Roche, TRIS Pharmaceuticals, Octapharma, and GW Pharmaceuticals, expert witness income from Lannett, and research contracts with Roche, Octapharma, and GW Pharmaceuticals for research unrelated to this manuscript. The other authors report no conflicts of interest.

### Competing Interest Statement

P.M.T. reports a research grant from Biogen, Inc. for research unrelated to this manuscript. J.T.M. reports consultant income from Roche, TRIS Pharmaceuticals, Octapharma, and GW Pharmaceuticals, expert witness income from Lannett, and research contracts with Roche, Octapharma, and GW Pharmaceuticals for research unrelated to this manuscript. The other authors report no conflicts of interest.

