## Supplemental Material for "White matter microstructure shows sex differences in late childhood: Evidence from 6,797 children"

### Sex differences in advanced measures of white matter microstructure in 6,797 children

#### *Supplementary Information*

#### Table S1. White matter fiber tract ROIs labeled by AtlasTrack.

| Abbreviation | Fiber Name | Category | Connected Brain Regions |
|---|---|---|---|
| ATR | Anterior thalamic radiations | Projection (thalamic) | Thalamus & frontal lobe |
| CC | Corpus callosum | Commissural | Left cortex & right cortex |
| CGC | Cingulum (cingulate) | Limbic | Cingulate gyrus & entorhinal cortex (cingulate portion) |
| CGH | Cingulum (parahippocampal) | Limbic | Cingulate gyrus & entorhinal cortex (parahippocampal portion) |
| CST | Corticospinal/pyramidal tract | Projection (sensorimotor) | Motor cortex & spinal cord |
| Fmaj | Forceps major | Commissural | Left occipital cortex & right occipital cortex |
| Fmin | Forceps minor | Commissural | Left prefrontal cortex & right prefrontal cortex |
| fSCS | Superior corticostriate (frontal cortex) | Projection (striatal) | Superior frontal cortex & striatum |
| FX | Fornix | Limbic | Hippocampus & mammilary nuclei of hypothalamus |
| FXcut | Fornix (excluding fimbria) | Limbic | Hippocampus & mammilary nuclei of hypothalamus (excluding fimbria) |
| IFO | Inferior fronto-occipital fasciculus | Association | Occipital lobe & frontal lobe |
| IFSFC | Inferior frontal superior frontal cortex | Association | Inferior frontal cortex & superior frontal cortex |
| ILF | Inferior longitudinal fasciculus | Association | Occipital lobe & temporal lobe |
| pSCS | Superior corticostriate (parietal cortex) | Projection (striatal) | Superior parietal cortex & striatum |
| pSLF | Superior longitudinal fasiculus (parietal) | Association | Parietal lobe & frontal lobe |
| SCS | Superior corticostriate | Projection (striatal) | Superior cortex & striatum |
| SIFC | Striatal inferior frontal cortex | Projection (striatal) | Inferior frontal cortex & striatum |
| SLF | Superior longitudinal fasciculus | Association | Temporal and parietal lobes & frontal lobe |
| tSLF | Superior longitudinal fasiculus (temporal) | Association | Temporal lobe & frontal lobe |
| UNC | Uncinate fasciculus | Association | Inferior frontal lobe & anterior temporal lobe |

#### Table S2. Regression analyses for sex differences in DTI-FA.

| | Discovery Cohort | | | | Replication Cohort | | |
|---|---|---|---|---|---|---|---|
| | **Std. Beta** | **SE** | **P** | **FDR** | **Std. Beta** | **SE** | **P** |
| All Fibers | 0.077 | 0.033 | 0.018 | 0.026 | 0.068 | 0.032 | 0.036 |
| ATR | -0.077 | 0.029 | 0.008 | 0.014 | -0.127 | 0.029 | 9.9E-06 |
| CC | 0.064 | 0.033 | 0.057 | 0.070 | 0.062 | 0.033 | 0.063 |
| CGC | -0.206 | 0.033 | 5.2E-10 | 5.4E-09 | -0.187 | 0.033 | 1.8E-08 |
| CGH | -0.076 | 0.032 | 0.017 | 0.026 | -0.087 | 0.032 | 0.007 |
| CST | 0.046 | 0.033 | 0.155 | 0.181 | 0.001 | 0.032 | 0.976 |
| Fmaj | 0.024 | 0.033 | 0.468 | 0.468 | -0.059 | 0.033 | 0.077 |
| Fmin | 0.066 | 0.032 | 0.041 | 0.054 | 0.091 | 0.033 | 0.005 |
| fSCS | 0.079 | 0.029 | 0.005 | 0.011 | 0.086 | 0.029 | 0.003 |
| FX | -0.136 | 0.032 | 1.8E-05 | 7.4E-05 | -0.203 | 0.031 | 9.5E-11 |
| FXcut | -0.099 | 0.031 | 0.002 | 0.004 | -0.174 | 0.031 | 2.1E-08 |
| IFO | 0.250 | 0.033 | 2.7E-14 | 5.7E-13 | 0.273 | 0.033 | 7.3E-17 |
| IFSFC | 0.182 | 0.032 | 2.0E-08 | 1.4E-07 | 0.168 | 0.032 | 2.1E-07 |
| ILF | 0.109 | 0.033 | 0.001 | 0.003 | 0.097 | 0.033 | 0.003 |
| pSCS | 0.134 | 0.029 | 6.1E-06 | 3.2E-05 | 0.116 | 0.03 | 9.8E-05 |
| pSLF | 0.035 | 0.033 | 0.297 | 0.311 | 0.041 | 0.033 | 0.219 |
| SCS | 0.103 | 0.029 | 3.4E-04 | 0.001 | 0.098 | 0.029 | 0.001 |
| SIFC | 0.093 | 0.033 | 0.005 | 0.011 | 0.149 | 0.033 | 7.0E-06 |
| SLF | 0.041 | 0.033 | 0.221 | 0.244 | 0.050 | 0.033 | 0.136 |
| tSLF | 0.089 | 0.033 | 0.008 | 0.014 | 0.103 | 0.033 | 0.002 |
| UNC | 0.083 | 0.034 | 0.015 | 0.024 | 0.098 | 0.034 | 0.004 |

#### Table S3. Regression analyses for sex differences in DTI-MD.

| | Discovery Cohort | | | | Replication Cohort | | |
|---|---|---|---|---|---|---|---|
| | **Std. Beta** | **SE** | **P** | **FDR** | **Std. Beta** | **SE** | **P** |
| All Fibers | -0.344 | 0.033 | 1.9E-25 | 6.8E-25 | -0.268 | 0.033 | 4.0E-16 |
| ATR | -0.258 | 0.032 | 2.8E-15 | 4.8E-15 | -0.166 | 0.033 | 3.9E-07 |
| CC | -0.282 | 0.033 | 1.9E-17 | 3.5E-17 | -0.190 | 0.033 | 8.0E-09 |
| CGC | -0.135 | 0.032 | 2.1E-05 | 2.4E-05 | -0.098 | 0.032 | 0.002 |
| CGH | -0.322 | 0.032 | 3.2E-23 | 7.5E-23 | -0.256 | 0.032 | 3.2E-15 |
| CST | -0.200 | 0.031 | 1.8E-10 | 2.3E-10 | -0.142 | 0.031 | 7.2E-06 |
| Fmaj | -0.171 | 0.033 | 2.2E-07 | 2.8E-07 | -0.066 | 0.033 | 0.044 |
| Fmin | -0.141 | 0.031 | 7.6E-06 | 8.8E-06 | -0.096 | 0.031 | 0.002 |
| fSCS | -0.325 | 0.031 | 7.2E-26 | 3.0E-25 | -0.314 | 0.030 | 1.4E-24 |
| FX | -0.079 | 0.034 | 0.020 | 0.020 | -0.004 | 0.034 | 0.899 |
| FXcut | -0.137 | 0.034 | 6.3E-05 | 0.001 | -0.058 | 0.034 | 0.087 |
| IFO | -0.489 | 0.032 | 1.4E-49 | 2.8E-48 | -0.428 | 0.033 | 1.4E-38 |
| IFSFC | -0.218 | 0.031 | 1.3E-12 | 2.0E-12 | -0.209 | 0.031 | 1.0E-11 |
| ILF | -0.417 | 0.033 | 3.6E-36 | 3.8E-35 | -0.419 | 0.033 | 4.4E-37 |
| pSCS | -0.304 | 0.031 | 3.0E-22 | 6.3E-22 | -0.280 | 0.031 | 4.5E-19 |
| pSLF | -0.323 | 0.031 | 4.1E-25 | 1.2E-24 | -0.296 | 0.031 | 2.6E-21 |
| SCS | -0.316 | 0.031 | 1.6E-24 | 4.1E-24 | -0.291 | 0.031 | 3.7E-21 |
| SIFC | -0.230 | 0.031 | 7.7E-14 | 1.2E-13 | -0.175 | 0.031 | 1.5E-08 |
| SLF | -0.343 | 0.032 | 6.3E-27 | 3.3E-26 | -0.323 | 0.032 | 4.2E-24 |
| tSLF | -0.357 | 0.032 | 4.5E-28 | 3.1E-27 | -0.341 | 0.032 | 8.8E-26 |
| UNC | -0.215 | 0.032 | 3.2E-11 | 4.5E-11 | -0.177 | 0.032 | 3.8E-08 |

##### Table S4. Regression analyses for sex differences in DTI-AD.

| | Discovery Cohort | | | | Replication Cohort | | |
|---|---|---|---|---|---|---|---|
| | **Std. Beta** | **SE** | **P** | **FDR** | **Std. Beta** | **SE** | **P** |
| All Fibers | -0.368 | 0.032 | 5.1E-30 | 2.2E-29 | -0.279 | 0.032 | 4.6E-18 |
| ATR | -0.289 | 0.031 | 5.4E-20 | 1.4E-19 | -0.251 | 0.031 | 2.0E-15 |
| CC | -0.309 | 0.032 | 1.5E-21 | 4.4E-21 | -0.196 | 0.032 | 1.2E-09 |
| CGC | -0.295 | 0.033 | 4.8E-19 | 1.1E-18 | -0.240 | 0.033 | 4.9E-13 |
| CGH | -0.380 | 0.031 | 1.6E-33 | 1.1E-32 | -0.330 | 0.031 | 1.5E-25 |
| CST | -0.147 | 0.031 | 1.8E-06 | 2.4E-06 | -0.122 | 0.030 | 6.4E-05 |
| Fmaj | -0.202 | 0.033 | 1.1E-09 | 1.8E-09 | -0.153 | 0.033 | 4.3E-06 |
| Fmin | -0.121 | 0.032 | 1.6E-04 | 1.7E-04 | -0.039 | 0.032 | 0.229 |
| fSCS | -0.240 | 0.033 | 6.1E-13 | 1.2E-12 | -0.219 | 0.033 | 3.8E-11 |
| FX | -0.142 | 0.034 | 2.5E-05 | 2.9E-05 | -0.106 | 0.034 | 1.7E-03 |
| FXcut | -0.169 | 0.034 | 7.4E-07 | 1.0E-06 | -0.119 | 0.034 | 4.3E-04 |
| IFO | -0.279 | 0.032 | 3.1E-18 | 6.5E-18 | -0.217 | 0.032 | 1.4E-11 |
| IFSFC | -0.118 | 0.031 | 1.6E-04 | 1.7E-04 | -0.116 | 0.031 | 1.9E-04 |
| ILF | -0.352 | 0.033 | 1.7E-25 | 5.8E-25 | -0.365 | 0.033 | 7.2E-28 |
| pSCS | -0.169 | 0.033 | 4.1E-07 | 6.2E-07 | -0.169 | 0.033 | 4.0E-07 |
| pSLF | -0.362 | 0.029 | 3.7E-34 | 3.8E-33 | -0.341 | 0.030 | 3.1E-30 |
| SCS | -0.212 | 0.033 | 2.3E-10 | 4.1E-10 | -0.194 | 0.033 | 5.0E-09 |
| SIFC | -0.098 | 0.029 | 5.7E-04 | 5.7E-04 | -0.010 | 0.029 | 0.714 |
| SLF | -0.387 | 0.030 | 4.4E-37 | 9.3E-36 | -0.374 | 0.030 | 2.4E-34 |
| tSLF | -0.369 | 0.031 | 8.6E-32 | 4.5E-31 | -0.352 | 0.031 | 1.2E-28 |
| UNC | -0.141 | 0.032 | 1.0E-05 | 1.2E-05 | -0.098 | 0.032 | 0.002 |

**Table S5. Regression analyses for sex differences in DTI-RD.**

| | Discovery Cohort | | | | Replication Cohort | | |
|---|---|---|---|---|---|---|---|
| | **Std. Beta** | **SE** | **P** | **FDR** | **Std. Beta** | **SE** | **P** |
| All Fibers | -0.249 | 0.033 | 5.5E-14 | 1.6E-13 | -0.198 | 0.033 | 2.2E-09 |
| ATR | -0.122 | 0.032 | 0.001 | 1.9E-04 | -0.032 | 0.032 | 0.310 |
| CC | -0.197 | 0.033 | 3.9E-09 | 6.8E-09 | -0.141 | 0.033 | 2.5E-05 |
| CGC | 0.089 | 0.032 | 0.006 | 0.006 | 0.094 | 0.032 | 0.004 |
| CGH | -0.153 | 0.033 | 3.5E-06 | 5.3E-06 | -0.108 | 0.033 | 0.001 |
| CST | -0.114 | 0.033 | 0.001 | 0.001 | -0.055 | 0.033 | 0.099 |
| Fmaj | -0.104 | 0.033 | 0.002 | 0.002 | 0.006 | 0.033 | 0.861 |
| Fmin | -0.116 | 0.031 | 2.3E-04 | 3.0E-04 | -0.105 | 0.031 | 0.001 |
| fSCS | -0.229 | 0.029 | 1.5E-15 | 7.8E-15 | -0.230 | 0.029 | 9.4E-16 |
| FX | -0.022 | 0.034 | 0.523 | 0.523 | 0.071 | 0.034 | 0.034 |
| FXcut | -0.099 | 0.034 | 0.004 | 0.004 | -0.006 | 0.034 | 0.864 |
| IFO | -0.433 | 0.033 | 2.5E-39 | 5.2E-38 | -0.404 | 0.033 | 1.0E-34 |
| IFSFC | -0.232 | 0.031 | 1.7E-13 | 4.0E-13 | -0.220 | 0.031 | 2.0E-12 |
| ILF | -0.309 | 0.033 | 9.0E-21 | 9.4E-20 | -0.301 | 0.033 | 5.9E-20 |
| pSCS | -0.236 | 0.030 | 4.6E-15 | 1.9E-14 | -0.214 | 0.030 | 1.6E-12 |
| pSLF | -0.211 | 0.033 | 1.2E-10 | 2.4E-10 | -0.192 | 0.033 | 4.4E-09 |
| SCS | -0.236 | 0.029 | 6.5E-16 | 4.5E-15 | -0.222 | 0.029 | 4.1E-14 |
| SIFC | -0.250 | 0.033 | 4.7E-14 | 1.6E-13 | -0.239 | 0.033 | 5.0E-13 |
| SLF | -0.217 | 0.033 | 5.5E-11 | 1.2E-10 | -0.203 | 0.033 | 8.1E-10 |
| tSLF | -0.245 | 0.033 | 1.6E-13 | 4.0E-13 | -0.236 | 0.033 | 1.1E-12 |
| UNC | -0.184 | 0.034 | 4.3E-08 | 6.9E-08 | -0.166 | 0.033 | 6.2E-07 |

#### Table S6. Regression analyses for sex differences in RSI-N0.

| | Discovery Cohort | | | | Replication Cohort | | |
|---|---|---|---|---|---|---|---|
| | Std. Beta | SE | P | FDR | Std. Beta | SE | P |
| All Fibers | 0.278 | 0.033 | 4.2E-17 | 8.0E-17 | 0.248 | 0.032 | 2.0E-14 |
| ATR | 0.136 | 0.033 | 3.1E-05 | 4.0E-05 | 0.076 | 0.032 | 0.017 |
| CC | 0.227 | 0.033 | 3.5E-12 | 5.6E-12 | 0.193 | 0.032 | 2.2E-09 |
| CGC | 0.055 | 0.032 | 0.084 | 0.093 | 0.036 | 0.032 | 0.252 |
| CGH | 0.296 | 0.031 | 1.7E-21 | 4.1E-21 | 0.242 | 0.031 | 5.2E-15 |
| CST | 0.254 | 0.031 | 1.1E-16 | 2.0E-16 | 0.251 | 0.03 | 1.1E-16 |
| Fmaj | 0.293 | 0.033 | 1.1E-18 | 2.4E-18 | 0.196 | 0.033 | 2.7E-09 |
| Fmin | 0.047 | 0.032 | 0.135 | 0.142 | 0.044 | 0.032 | 0.161 |
| fSCS | 0.380 | 0.033 | 2.7E-30 | 2.8E-29 | 0.405 | 0.032 | 5.6E-36 |
| FX | 0.046 | 0.033 | 0.165 | 0.165 | -0.042 | 0.033 | 0.205 |
| FXcut | 0.097 | 0.034 | 0.004 | 0.005 | -3.0E-04 | 0.034 | 0.993 |
| IFO | 0.450 | 0.033 | 4.2E-42 | 8.8E-41 | 0.419 | 0.032 | 2.3E-37 |
| IFSFC | 0.172 | 0.033 | 1.5E-07 | 2.2E-07 | 0.190 | 0.032 | 2.9E-09 |
| ILF | 0.378 | 0.033 | 2.3E-29 | 1.2E-28 | 0.386 | 0.033 | 5.3E-31 |
| pSCS | 0.344 | 0.033 | 5.5E-25 | 1.5E-24 | 0.334 | 0.033 | 2.5E-24 |
| pSLF | 0.340 | 0.032 | 1.6E-25 | 4.8E-25 | 0.317 | 0.032 | 5.6E-23 |
| SCS | 0.367 | 0.033 | 2.4E-28 | 8.5E-28 | 0.361 | 0.032 | 1.2E-28 |
| SIFC | 0.161 | 0.033 | 9.9E-07 | 1.4E-06 | 0.132 | 0.032 | 4.5E-05 |
| SLF | 0.366 | 0.033 | 8.6E-29 | 3.6E-28 | 0.352 | 0.032 | 2.3E-27 |
| tSLF | 0.374 | 0.033 | 1.6E-29 | 1.1E-28 | 0.362 | 0.032 | 2.1E-28 |
| UNC | 0.103 | 0.033 | 0.002 | 0.002 | 0.086 | 0.033 | 0.009 |

#### Table S7. Regression analyses for sex differences in RSI-ND.

| | Discovery Cohort | | | | Replication Cohort | | |
|---|---|---|---|---|---|---|---|
| | **Std. Beta** | **SE** | **P** | **FDR** | **Std. Beta** | **SE** | **P** |
| All Fibers | 0.094 | 0.032 | 0.004 | 0.009 | 0.078 | 0.032 | 0.017 |
| ATR | -0.049 | 0.032 | 0.123 | 0.185 | -0.099 | 0.032 | 0.002 |
| CC | 0.080 | 0.031 | 0.010 | 0.024 | 0.071 | 0.031 | 0.023 |
| CGC | -0.171 | 0.033 | 1.9E-07 | 1.3E-06 | -0.160 | 0.033 | 1.3E-06 |
| CGH | 0.019 | 0.033 | 0.563 | 0.591 | -0.013 | 0.033 | 0.690 |
| CST | 0.036 | 0.033 | 0.274 | 0.302 | 0.004 | 0.033 | 0.902 |
| Fmaj | 0.057 | 0.032 | 0.073 | 0.127 | -0.043 | 0.032 | 0.174 |
| Fmin | 0.036 | 0.029 | 0.215 | 0.250 | 0.054 | 0.029 | 0.064 |
| fSCS | 0.114 | 0.031 | 2.4E-04 | 0.001 | 0.144 | 0.031 | 3.2E-06 |
| FX | -0.042 | 0.033 | 0.209 | 0.250 | -0.147 | 0.033 | 8.8E-06 |
| FXcut | -0.004 | 0.033 | 0.909 | 0.909 | -0.125 | 0.033 | 1.7E-04 |
| IFO | 0.272 | 0.030 | 9.3E-20 | 1.9E-18 | 0.294 | 0.030 | 1.6E-22 |
| IFSFC | 0.145 | 0.033 | 9.7E-06 | 4.1E-05 | 0.143 | 0.033 | 1.3E-05 |
| ILF | 0.177 | 0.033 | 1.3E-07 | 1.3E-06 | 0.147 | 0.033 | 1.0E-05 |
| pSCS | 0.150 | 0.031 | 1.4E-06 | 7.2E-06 | 0.142 | 0.031 | 5.4E-06 |
| pSLF | 0.048 | 0.032 | 0.142 | 0.187 | 0.023 | 0.033 | 0.484 |
| SCS | 0.126 | 0.031 | 3.6E-05 | 1.3E-04 | 0.134 | 0.030 | 1.1E-05 |
| SIFC | 0.049 | 0.030 | 0.104 | 0.169 | 0.093 | 0.030 | 0.002 |
| SLF | 0.048 | 0.032 | 0.137 | 0.187 | 0.030 | 0.033 | 0.353 |
| tSLF | 0.077 | 0.032 | 0.018 | 0.038 | 0.073 | 0.033 | 0.026 |
| UNC | 0.063 | 0.032 | 0.054 | 0.102 | 0.065 | 0.033 | 0.046 |

#### Table S8. Regression analyses for sex differences in RSI-NT.

| | Discovery Cohort | | | | Replication Cohort | | |
|---|---|---|---|---|---|---|---|
| | Std. Beta | SE | P | FDR | Std. Beta | SE | P |
| All Fibers | 0.159 | 0.033 | 1.6E-06 | 4.2E-06 | 0.140 | 0.033 | 2.0E-05 |
| ATR | 0.013 | 0.033 | 0.696 | 0.696 | -0.046 | 0.032 | 0.153 |
| CC | 0.121 | 0.032 | 1.5E-04 | 2.7E-04 | 0.110 | 0.032 | 0.001 |
| CGC | -0.144 | 0.033 | 1.5E-05 | 2.8E-05 | -0.141 | 0.033 | 2.1E-05 |
| CGH | 0.089 | 0.033 | 0.006 | 0.008 | 0.049 | 0.033 | 0.140 |
| CST | 0.111 | 0.033 | 0.001 | 0.001 | 0.086 | 0.033 | 0.009 |
| Fmaj | 0.116 | 0.032 | 2.6E-04 | 4.2E-04 | 0.009 | 0.032 | 0.775 |
| Fmin | 0.040 | 0.030 | 0.180 | 0.210 | 0.056 | 0.030 | 0.062 |
| fSCS | 0.241 | 0.032 | 3.8E-14 | 4.0E-13 | 0.274 | 0.031 | 3.7E-18 |
| FX | -0.019 | 0.033 | 0.563 | 0.591 | -0.126 | 0.033 | 1.5E-04 |
| FXcut | 0.029 | 0.034 | 0.393 | 0.434 | -0.097 | 0.034 | 0.004 |
| IFO | 0.335 | 0.031 | 2.9E-27 | 6.1E-26 | 0.346 | 0.031 | 6.3E-29 |
| IFSFC | 0.171 | 0.033 | 3.2E-07 | 9.6E-07 | 0.177 | 0.033 | 1.0E-07 |
| ILF | 0.249 | 0.033 | 1.4E-13 | 5.7E-13 | 0.226 | 0.033 | 1.2E-11 |
| pSCS | 0.239 | 0.032 | 8.1E-14 | 4.3E-13 | 0.230 | 0.032 | 5.0E-13 |
| pSLF | 0.152 | 0.033 | 5.8E-06 | 1.2E-05 | 0.127 | 0.033 | 1.4E-04 |
| SCS | 0.235 | 0.031 | 8.0E-14 | 4.3E-13 | 0.241 | 0.031 | 1.5E-14 |
| SIFC | 0.098 | 0.031 | 0.002 | 0.003 | 0.123 | 0.031 | 9.2E-05 |
| SLF | 0.158 | 0.033 | 2.1E-06 | 4.9E-06 | 0.142 | 0.033 | 1.9E-05 |
| tSLF | 0.181 | 0.033 | 5.9E-08 | 2.1E-07 | 0.177 | 0.033 | 1.1E-07 |
| UNC | 0.075 | 0.033 | 0.023 | 0.028 | 0.073 | 0.033 | 0.027 |

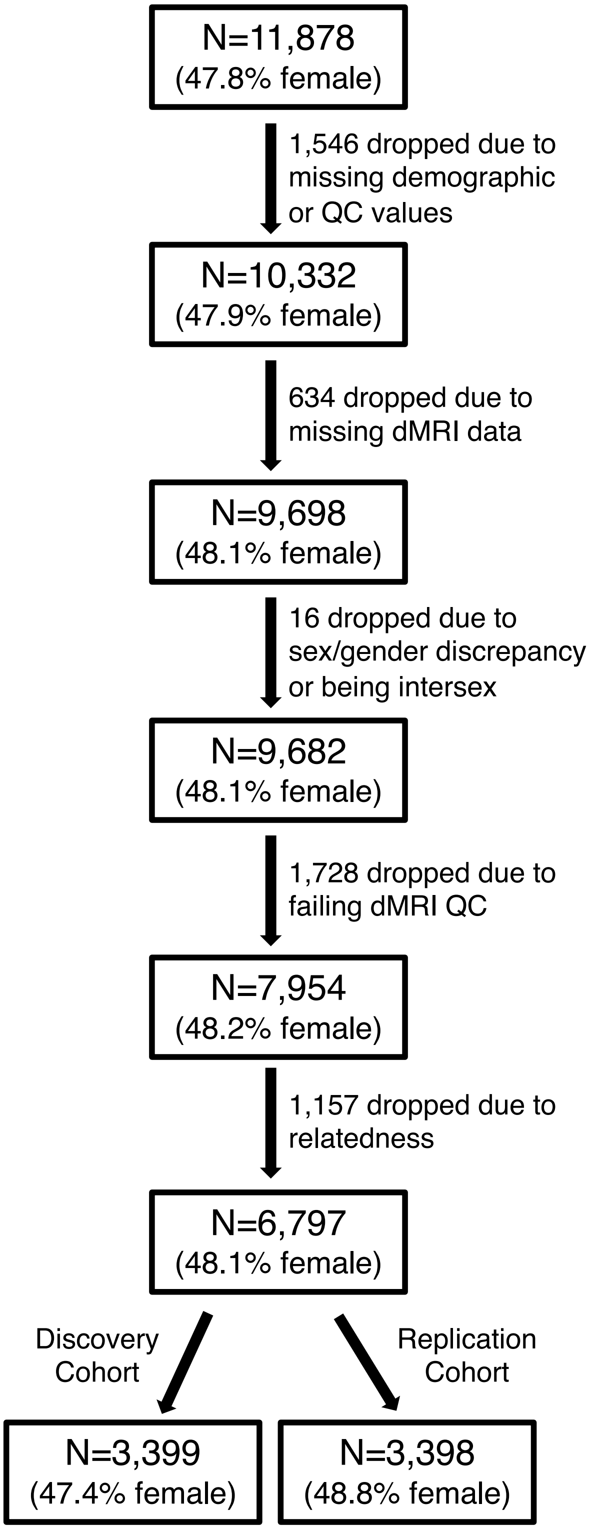

**Figure S1. Participant flowchart.** Subjects were only included in the current analyses if they had complete nuisance covariate data (age, household income, parental education, race, ethnicity, MRI scanner), dMRI quality control scores, and complete dMRI data that also passed quality control. Intersex participants, or participants whose biological sex assigned at birth did not align with their parent-report gender identity, were excluded from analyses. Children were additionally excluded if they were the sibling of another subject in the study, with the retained sibling selected randomly. Our final sample was split into a discovery cohort and a replication cohort to demonstrate the reproducibility of all findings. dMRI = diffusion-weighted MRI.

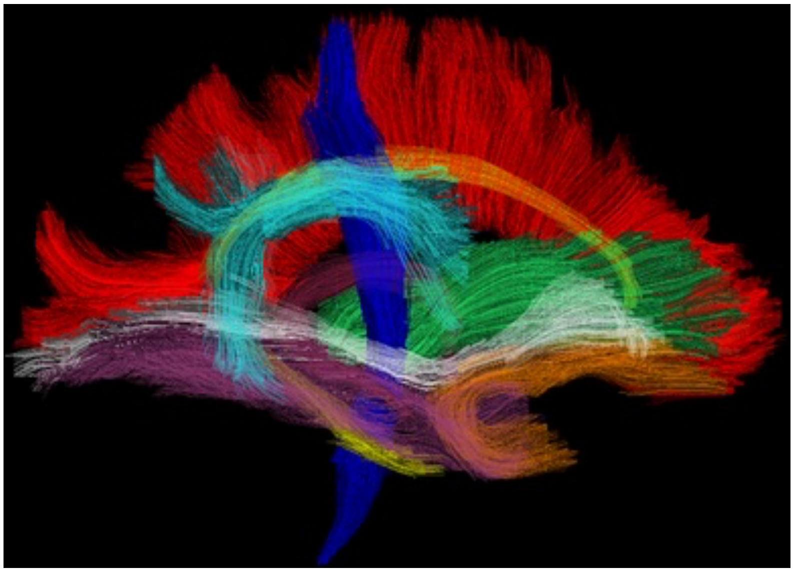

**Figure S2. Example white matter fiber tracts from AtlasTrack.** Anterior thalamic radiation (ATR) = green; corpus callosum (CC) = red; cingulate and parahippocampal cingulum (CGC, CGH) = yellow; *forceps major* (Fmaj) = red; *forceps minor* (Fmin) = red; fornix (FX) = magenta; inferior fronto-occipital fasciculus (IFO) = white; inferior longitudinal fasciculus (ILF) = purple; full, parietal, and temporal superior longitudinal fasciculus (SLF, pSLF, tSLF) = cyan; uncinate fasciculus (UNC) = orange. Image reproduced from the Neuroimaging Tools & Resource Collaboratory (NITRC) distribution of AtlasTrack (https://www.nitrc.org/projects/atlastrack/); see below for complete copyright information.

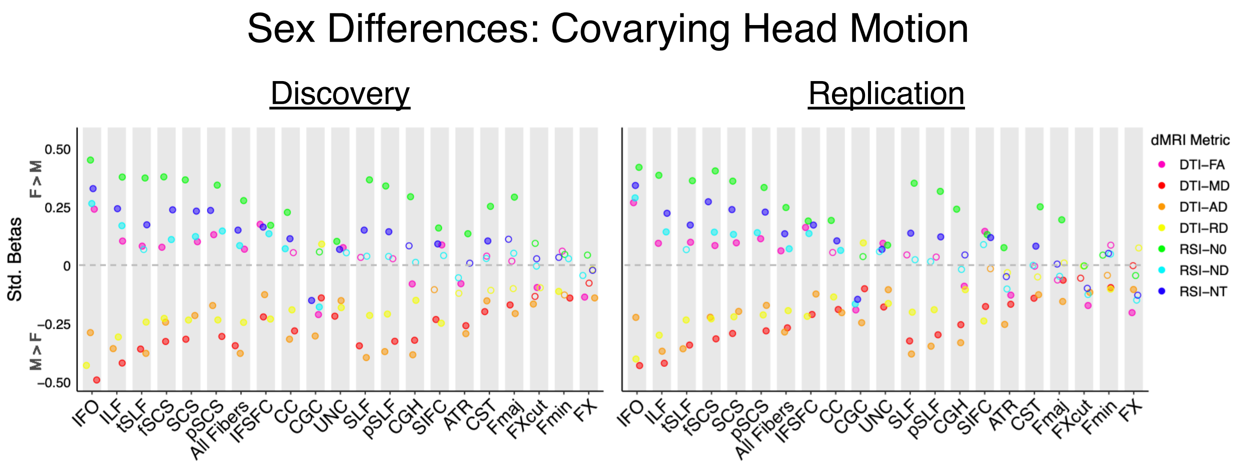

**Figure S3. Sex differences in white matter microstructure when controlling for head motion**. The effect size, significance, and replicability of sex differences in DTI and RSI metrics are depicted separately for the discovery cohort (left) and replication cohort (right) for bilateral white matter tract ROIs. Positive standardized betas indicate Female > Male, and negative standardized betas indicate Male > Female. DTI metrics are depicted in warm colors, and RSI metrics in cool colors. Filled circles indicate the association was both significant in the discovery cohort after FDR correction across the number of ROIs ($q$<0.05) and demonstrated $p$<0.05 in the replication cohort. *Diffusion-weighted MRI abbreviations*: ROI = region of interest, dMRI = diffusion-weighted MRI, DTI = diffusion tensor imaging, FA = fractional anisotropy, MD = mean diffusivity, AD = axial diffusivity, RD = radial diffusivity, RSI = restriction spectrum imaging, N0 = normalized isotropic, ND = normalized directional, NT = normalized total. *White matter tract ROI abbreviations*: ATR = anterior thalamic radiation, CC = corpus callosum, CGC = cingulum (cingulate), CGH = cingulum (parahippocampal), CST = corticospinal/pyramidal tract, Fmaj = forceps major, Fmin = forceps minor, fSCS = superior corticostriate (frontal cortex), FX = fornix, FXcut = fornix (excluding fimbria), IFO = inferior fronto-occipital fasciculus, IFSFC = inferior frontal superior frontal cortex, ILF = inferior longitudinal fasciculus, pSCS = superior corticostriate (parietal cortex), pSLF = superior longitudinal fasciculus (parietal), SCS = superior corticostriate, SIFC = striatal inferior frontal cortex, SLF = superior longitudinal fasciculus, tSLF = superior longitudinal fasciculus (temporal), UNC = uncinate fasciculus.

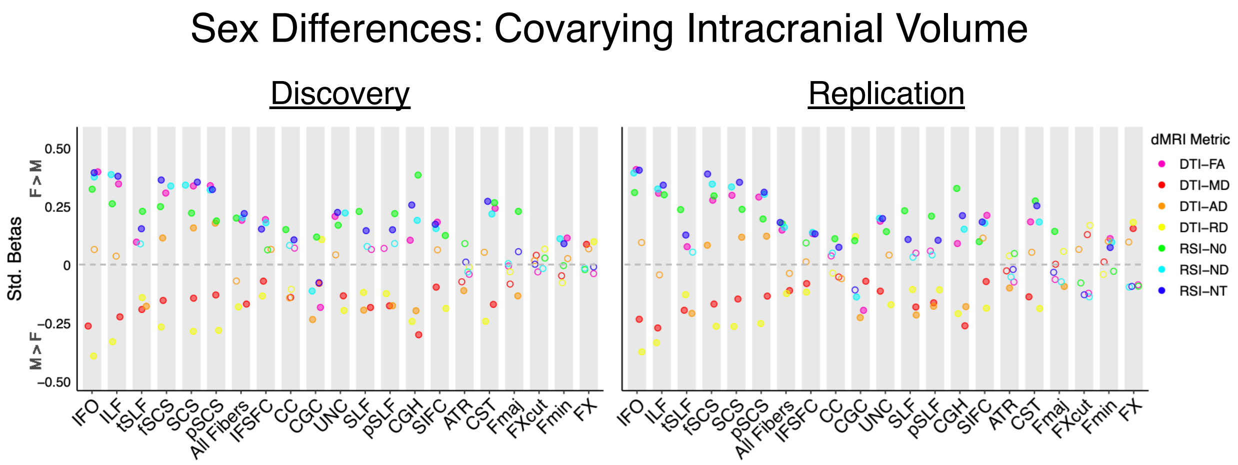

**Figure S4. Sex differences in white matter microstructure when controlling for intracranial volume**. The effect size, significance, and replicability of sex differences in DTI and RSI metrics are depicted separately for the discovery cohort (left) and replication cohort (right) for bilateral white matter tract ROIs. Positive standardized betas indicate Female > Male, and negative standardized betas indicate Male > Female. DTI metrics are depicted in warm colors, and RSI metrics in cool colors. Filled circles indicate the association was both significant in the discovery cohort after FDR correction across the number of ROIs ($q<0.05$) and demonstrated $p<0.05$ in the replication cohort. *Diffusion-weighted MRI abbreviations*: ROI = region of interest, dMRI = diffusion-weighted MRI, DTI = diffusion tensor imaging, FA = fractional anisotropy, MD = mean diffusivity, AD = axial diffusivity, RD = radial diffusivity, RSI = restriction spectrum imaging, N0 = normalized isotropic, ND = normalized directional, NT = normalized total. *White matter tract ROI abbreviations*: ATR = anterior thalamic radiation, CC = corpus callosum, CGC = cingulum (cingulate), CGH = cingulum (parahippocampal), CST = corticospinal/pyramidal tract, Fmaj = forceps major, Fmin = forceps minor, fSCS = superior corticostriate (frontal cortex), FX = fornix, FXcut = fornix (excluding fimbria), IFO = inferior fronto-occipital fasciculus, IFSFC = inferior frontal superior frontal cortex, ILF = inferior longitudinal fasciculus, pSCS = superior corticostriate (parietal cortex), pSLF = superior longitudinal fasciculus (parietal), SCS = superior corticostriate, SIFC = striatal inferior frontal cortex, SLF = superior longitudinal fasciculus, tSLF = superior longitudinal fasciculus (temporal), UNC = uncinate fasciculus.

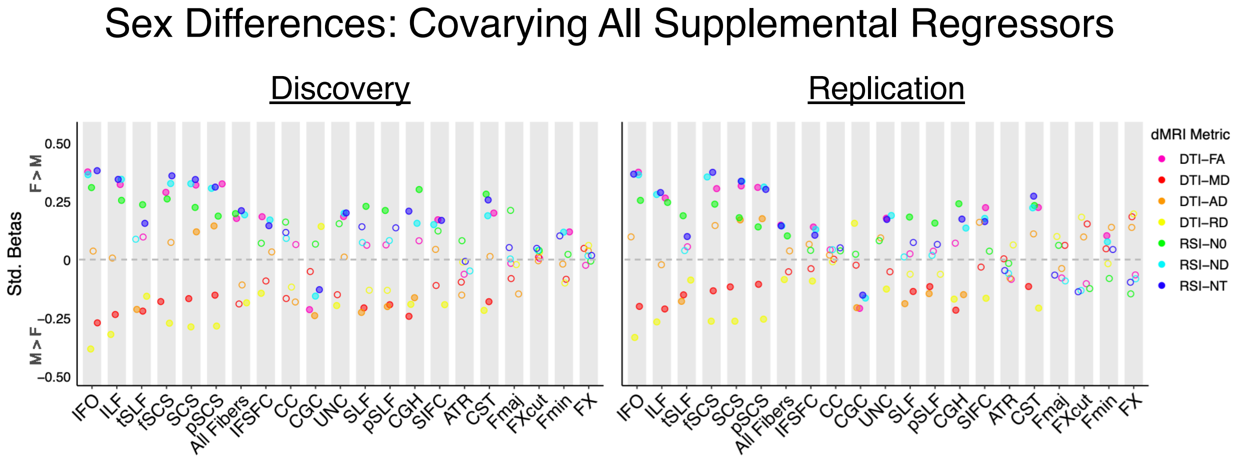

**Figure S5. Sex differences in white matter microstructure when controlling for all supplemental nuisance covariates in a single analysis (pubertal development, externalizing problems, internalizing problems, head motion, and intracranial volume)**. The effect size, significance, and replicability of sex differences in DTI and RSI metrics are depicted separately for the discovery cohort (left) and replication cohort (right) for bilateral white matter tract ROIs. Positive standardized betas indicate Female > Male, and negative standardized betas indicate Male > Female. DTI metrics are depicted in warm colors, and RSI metrics in cool colors. Filled circles indicate the association was both significant in the discovery cohort after FDR correction across the number of ROIs ($q<0.05$) and demonstrated $p<0.05$ in the replication cohort. *Diffusion-weighted MRI abbreviations*: ROI = region of interest, dMRI = diffusion-weighted MRI, DTI = diffusion tensor imaging, FA = fractional anisotropy, MD = mean diffusivity, AD = axial diffusivity, RD = radial diffusivity, RSI = restriction spectrum imaging, N0 = normalized isotropic, ND = normalized directional, NT = normalized total. *White matter tract ROI abbreviations*: ATR = anterior thalamic radiation, CC = corpus callosum, CGC = cingulum (cingulate), CGH = cingulum (parahippocampal), CST = corticospinal/pyramidal tract, Fmaj = forceps major, Fmin = forceps minor, fSCS = superior corticostriate (frontal cortex), FX = fornix, FXcut = fornix (excluding fimbria), IFO = inferior fronto-occipital fasciculus, IFSFC = inferior frontal superior frontal cortex, ILF = inferior longitudinal fasciculus, pSCS = superior corticostriate (parietal cortex), pSLF = superior longitudinal fasciculus (parietal), SCS = superior corticostriate, SIFC = striatal inferior frontal cortex, SLF = superior longitudinal fasciculus, tSLF = superior longitudinal fasciculus (temporal), UNC = uncinate fasciculus.

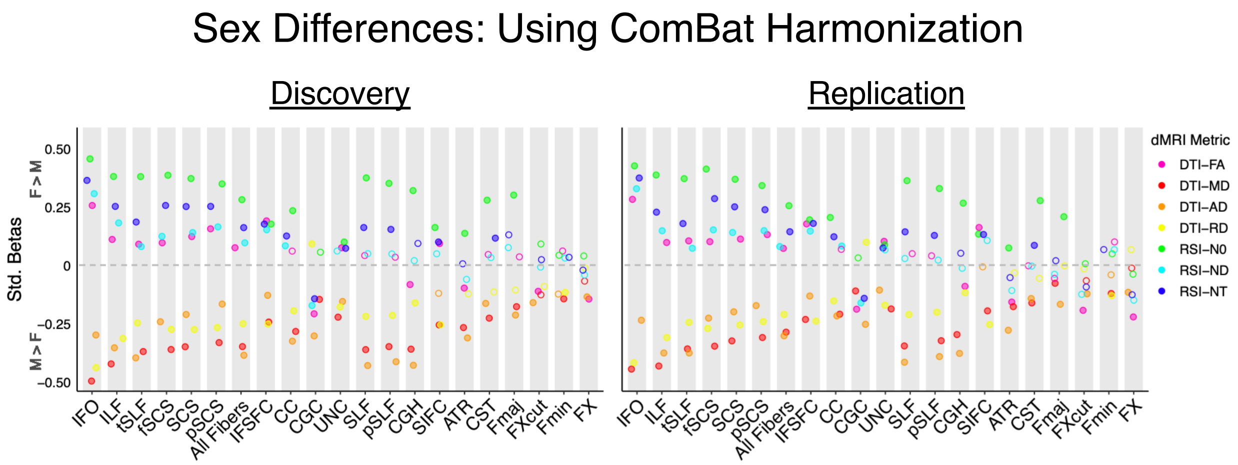

**Figure S6. Sex differences in white matter microstructure when using the advanced scanner harmonization approach, ComBat**. The effect size, significance, and replicability of sex differences in DTI and RSI metrics are depicted separately for the discovery cohort (left) and replication cohort (right) for bilateral white matter tract ROIs. Positive standardized betas indicate Female > Male, and negative standardized betas indicate Male > Female. DTI metrics are depicted in warm colors, and RSI metrics in cool colors. Filled circles indicate the association was both significant in the discovery cohort after FDR correction across the number of ROIs ($q$<0.05) and demonstrated $p$<0.05 in the replication cohort. *Diffusion-weighted MRI abbreviations*: ROI = region of interest, dMRI = diffusion-weighted MRI, DTI = diffusion tensor imaging, FA = fractional anisotropy, MD = mean diffusivity, AD = axial diffusivity, RD = radial diffusivity, RSI = restriction spectrum imaging, N0 = normalized isotropic, ND = normalized directional, NT = normalized total. *White matter tract ROI abbreviations*: ATR = anterior thalamic radiation, CC = corpus callosum, CGC = cingulum (cingulate), CGH = cingulum (parahippocampal), CST = corticospinal/pyramidal tract, Fmaj = forceps major, Fmin = forceps minor, fSCS = superior corticostriate (frontal cortex), FX = fornix, FXcut = fornix (excluding fimbria), IFO = inferior fronto-occipital fasciculus, IFSFC = inferior frontal superior frontal cortex, ILF = inferior longitudinal fasciculus, pSCS = superior corticostriate (parietal cortex), pSLF = superior longitudinal fasciculus (parietal), SCS = superior corticostriate, SIFC = striatal inferior frontal cortex, SLF = superior longitudinal fasciculus, tSLF = superior longitudinal fasciculus (temporal), UNC = uncinate fasciculus.

### Sex Differences: Combining Cohorts

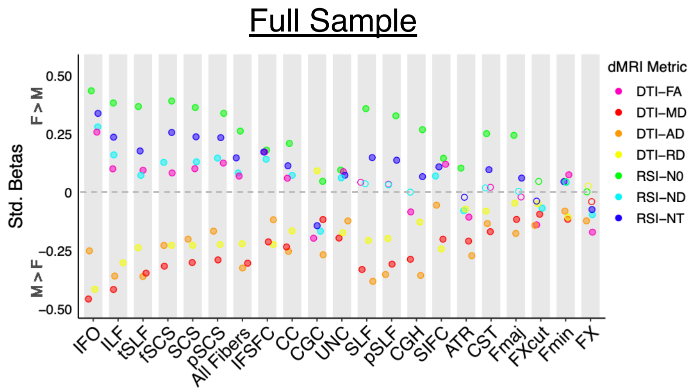

**Figure S7. Sex differences in white matter microstructure when combining discovery and replication cohorts into a single cohort**. The effect size, significance, and replicability of sex differences in DTI and RSI metrics are depicted for the full sample for bilateral white matter tract ROIs. Positive standardized betas indicate Female > Male, and negative standardized betas indicate Male > Female. DTI metrics are depicted in warm colors, and RSI metrics in cool colors. Filled circles indicate the association was significant after FDR correction across the number of ROIs ($q$<0.05). *Diffusion-weighted MRI abbreviations*: ROI = region of interest, dMRI = diffusion-weighted MRI, DTI = diffusion tensor imaging, FA = fractional anisotropy, MD = mean diffusivity, AD = axial diffusivity, RD = radial diffusivity, RSI = restriction spectrum imaging, N0 = normalized isotropic, ND = normalized directional, NT = normalized total. *White matter tract ROI abbreviations*: ATR = anterior thalamic radiation, CC = corpus callosum, CGC = cingulum (cingulate), CGH = cingulum (parahippocampal), CST = corticospinal/pyramidal tract, Fmaj = forceps major, Fmin = forceps minor, fSCS = superior corticostriate (frontal cortex), FX = fornix, FXcut = fornix (excluding fimbria), IFO = inferior fronto-occipital fasciculus, IFSFC = inferior frontal superior frontal cortex, ILF = inferior longitudinal fasciculus, pSCS = superior corticostriate (parietal cortex), pSLF = superior longitudinal fasciculus (parietal), SCS = superior corticostriate, SIFC = striatal inferior frontal cortex, SLF = superior longitudinal fasciculus, tSLF = superior longitudinal fasciculus (temporal), UNC = uncinate fasciculus.

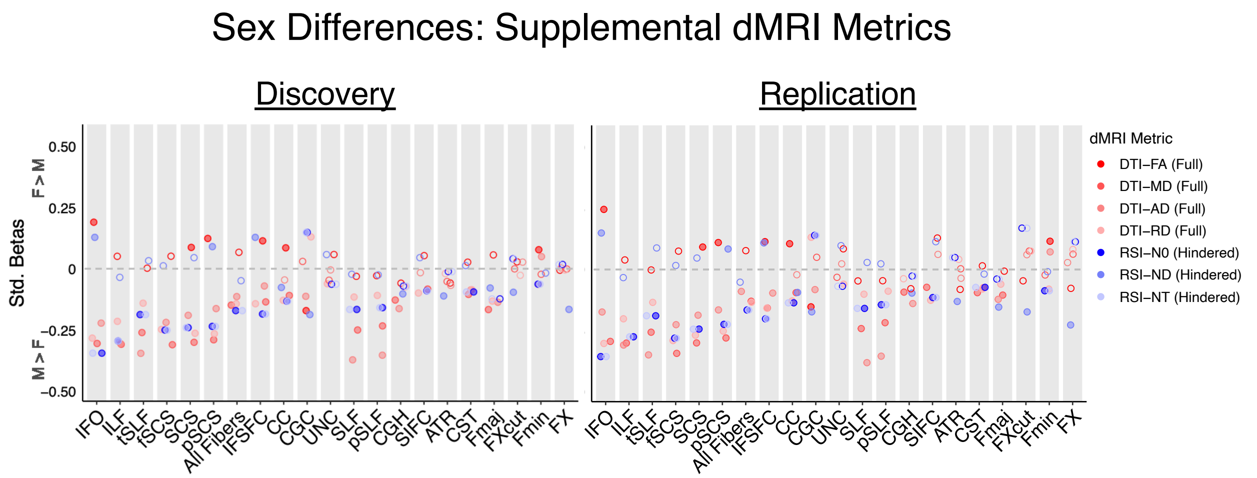

**Figure S8. Sex differences in white matter microstructure for supplemental diffusion-weighted MRI metrics.** The effect size, significance, and replicability of sex differences in full shell DTI and hindered (extracellular) RSI metrics are depicted separately for the discovery cohort (left) and replication cohort (right) for bilateral white matter tract ROIs. Note that the full shell DTI and hindered RSI metrics presented here represent different underlying diffusion properties than the inner shell DTI and restricted RSI metrics presented elsewhere in the manuscript; these supplemental results are thus not directly comparable to the primary findings focused on inner shell DTI and restricted RSI measures. Positive standardized betas indicate Female > Male, and negative standardized betas indicate Male > Female. Full shell DTI metrics are depicted in warm colors, and hindered RSI metrics in cool colors. Filled circles indicate the association was both significant in the discovery cohort after FDR correction across the number of ROIs ($q$<0.05) and demonstrated $p$<0.05 in the replication cohort. *Diffusion-weighted MRI abbreviations*: ROI = region of interest, dMRI = diffusion-weighted MRI, DTI = diffusion tensor imaging, FA = fractional anisotropy, MD = mean diffusivity, AD = axial diffusivity, RD = radial diffusivity, RSI = restriction spectrum imaging, N0 = normalized isotropic, ND = normalized directional, NT = normalized total. *White matter tract ROI abbreviations*: ATR = anterior thalamic radiation, CC = corpus callosum, CGC = cingulum (cingulate), CGH = cingulum (parahippocampal), CST = corticospinal/pyramidal tract, Fmaj = forceps major, Fmin = forceps minor, fSCS = superior corticostriate (frontal cortex), FX = fornix, FXcut = fornix (excluding fimbria), IFO = inferior fronto-occipital fasciculus, IFSFC = inferior frontal superior frontal cortex, ILF = inferior longitudinal fasciculus, pSCS = superior corticostriate (parietal cortex), pSLF = superior longitudinal fasciculus (parietal), SCS = superior corticostriate, SIFC = striatal inferior frontal cortex, SLF = superior longitudinal fasciculus, tSLF = superior longitudinal fasciculus (temporal), UNC = uncinate fasciculus.

#### Supplemental Text: Complete Copyright Information for AtlasTrack

Copyright Notice

This software is Copyright © 2020 The Regents of the University of California. All Rights Reserved. Permission to copy, modify, and distribute this software and its documentation for educational, research and non-profit purposes, without fee, and without a written agreement is hereby granted, provided that the above copyright notice, this paragraph and the following three paragraphs appear in all copies. Permission to make commercial use of this software may be obtained by contacting:

Office of Innovation and Commercialization

9500 Gilman Drive, Mail Code 0910

University of California

La Jolla, CA 92093-0910

(858) 534-5815

This software program and documentation are copyrighted by The Regents of the University of California. The software program and documentation are supplied “as is”, without any accompanying services from The Regents. The Regents does not warrant that the operation of the program will be uninterrupted or error-free. The end-user understands that the program was developed for research purposes and is advised not to rely exclusively on the program for any reason.

IN NO EVENT SHALL THE UNIVERSITY OF CALIFORNIA BE LIABLE TO ANY PARTY FOR DIRECT, INDIRECT, SPECIAL, INCIDENTAL, OR CONSEQUENTIAL DAMAGES, INCLUDING LOST PROFITS, ARISING OUT OF THE USE OF THIS SOFTWARE AND ITS DOCUMENTATION, EVEN IF THE UNIVERSITY OF CALIFORNIA HAS BEEN ADVISED OF THE POSSIBILITY OF SUCH DAMAGE. THE UNIVERSITY OF CALIFORNIA SPECIFICALLY DISCLAIMS ANY WARRANTIES, INCLUDING, BUT NOT LIMITED TO, THE IMPLIED WARRANTIES OF MERCHANTABILITY AND FITNESS FOR A PARTICULAR PURPOSE. THE SOFTWARE PROVIDED HEREUNDER IS ON AN “AS IS” BASIS, AND THE UNIVERSITY OF CALIFORNIA HAS NO OBLIGATIONS TO PROVIDE MAINTENANCE, SUPPORT, UPDATES, ENHANCEMENTS, OR MODIFICATIONS.
